# Prey speed up, predators slow down: non-consumptive effects on movement behavior of a ciliate predator-prey pair

**DOI:** 10.1101/2021.11.15.468607

**Authors:** Uriah Daugaard, Reinhard Furrer, Owen L. Petchey

**Affiliations:** Department of Evolutionary Biology and Environmental Studies, University of Zurich, Winterthurerstrasse 190, 8057 Zurich, Switzerland; Department of Mathematics and Computational Science, University of Zurich, Winterthurerstrasse 190, 8057 Zurich, Switzerland

**Keywords:** Predator-prey interactions, foraging, behavioral plasticity, protist, anti-predator defenses, movement ecology

## Abstract

Non-consumptive effects (NCEs) of predators on prey, such as induced defensive strategies, are frequently neglected in the analysis of predator-prey interactions. Yet these effects can have demographic impacts as strong as consumption. As a counterpart to NCEs, resource-availability effects (RAEs) can prompt changes in predators as well, e.g., in their foraging behavior. We studied NCEs and RAEs in the ciliate predator-prey pair *Didinium nasutum* and *Paramecium caudatum*. We examined the dependence of prey/predator swimming speed and body size on predator/prey presence. We also investigated prey spatial grouping behavior and the dependence of predator movement on local prey abundance. We collected individual movement and morphology data through videography of laboratory-based populations. We compared swimming speeds and body sizes based on their distributions. We used linear models to respectively quantify the effects of local prey abundance on predator displacements and of predator presence on prey grouping behavior. In the presence of prey, predator individuals swam more slowly, were bigger and made smaller displacements. Further, their displacements decreased with increasing local prey abundance. In contrast, in the presence of predators, proportionally more prey individuals showed a fast-swimming behavior and there was weak evidence for increased prey grouping. Trait changes entail energy expenditure shifts, which likely affect interspecific interactions and populations, as has been shown for NCEs. Less is known about the link between RAEs and demography, but it seems likely that the observed effects scale up to influence community and ecosystem stability, yet this remains largely unexplored.

**Significance Statement:** To maximize their fitness, organisms balance investment in foraging and avoiding being eaten. The behaviors of prey and predators are thus expected to depend on the presence and absence of each other and serve either to boost the chances of predation evasion or to increase predation success. Here we provide an example of the co-dependence of behaviors in the predator-prey pair *Didinium nasutum* and *Paramecium caudatum*. We show that the predator slows down and searches in smaller areas when prey are present, while the prey speeds up and possibly groups more as a response to the presence of predators. Such behavioral changes are likely to have a demographic and community impact that is not accounted for with common measures of predators-prey interactions.

## Introduction

Predation is an ubiquitous interspecific interaction and can contribute to the long-term coexistence of species (MacArthur 1984; Chesson 2000). Keystone species are known to maintain biodiversity (Paine 1969; Mills et al. 1993) while more recent results suggest that weak consumer-resource interactions can stabilize a food web containing multiple strong interactions (O’Gorman and Emmerson 2009; Kadoya et al. 2018).

In addition to consuming their prey, predators also have non-consumptive effects (NCEs, also known as trait-mediated interactions and as non-lethal effects). These include inducing defensive strategies and stress-induced changes in behavior (Lima 1998; Werner and Peacor 2003; Hermann and Landis 2017; MacLeod et al. 2018). For the prey, these changes can lead to a decreased energy intake, increased energy expenditure for the defense against predation, and decreased growth rates (e.g. Preisser et al. 2005; Clinchy et al. 2013). These effects scale up to have an impact on prey demography that can be at least as strong as the one caused by direct consumption (Preisser et al. 2005). NCEs are thus hypothesized to affect community stability (Anholt and Werner 1995; Peckarsky et al. 2008), with some evidence that this is the case (van Veen et al. 2005; Fill et al. 2012)

Similarly to the prey reacting to predator-presence, predators are expected to adapt their traits to the abundance of their prey. Here, we refer to this dependency as resource availability effects (RAEs), which can be seen as trait mediated interactions from the prey to the predator. For instance, predators can increase their foraging efforts when prey are scarce (Munk 1995; Ronconi and Burger 2008). In general, predator characteristics such as optimal patch use (e.g. McNamara 1982; Wajnberg et al. 2006) and optimal diet and foraging (O’brien et al. 1990; Sih and Christensen 2001) are likely to depend on prey abundance (i.e. resource availability). As an example of the prey density dependence of these processes, a simple Gaussian random walk is sufficient as a search strategy for the foraging of abundant prey, while a more efficient search strategy becomes increasingly important with decreasing resource availability (Humphries et al. 2010).

A predator-prey interaction is thus defined by the consumptive and the non-consumptive effects of the predator on the prey and by the resource availability effects of the prey on the predator. Yet, predator-prey interaction strengths are frequently calculated by only considering predator feeding rates as a function of prey abundance (i.e. the functional response Berlow et al. 2004; Rall et al. 2010; Sentis et al. 2012; Kalinoski and DeLong 2016; Daugaard et al. 2019). The functional response represents the mortality rate of the prey due to the consumption by the predator. Its product with the conversion efficiency is often used in models as the increase rate of the predator population (e.g. Yodzis 1994; Abrams and Ginzburg 2000). In other words, this approach does not account for any NCEs and RAEs impacting the growth rate of the prey (Van Buskirk and Yurewicz 1998; Preisser and Bolnick 2008; Sheriff et al. 2020) and the mortality rate and conversion efficiency of the predator (Bozer et al. 1996; Minter et al. 2011; Li and Montagnes 2015). To thoroughly characterize such a predator-prey pair, it is thus not enough to limit the analysis to the functional response (Lima 1998; Schmitz 2017).

Here we investigated the presence and characteristics of NCEs and RAEs in a microbial predator-prey pair of the ciliates *Didinium nasutum* (predator) and *Paramecium caudatum* (prey). Due to their favorable traits (e.g. short generation times and low maintenance) ciliates are frequently used in laboratory-based experiments for gaining insights into processes and patterns that are difficult to discern otherwise (Altermatt et al. 2015). Indeed, some of the earliest experiments about predator-prey dynamics were performed with *Didinium* and *Paramecium* (Luckinbill 1974). While simple (i.e., unicellular organisms without a nervous system), ciliates display various reactions to their environment such as bending, ciliary alteration, contractions and detachment (Jennings 1902; Dexter et al. 2019). Prey ciliate species have been observed to significantly reduce their swimming speed, increase their body width and to show additional defenses, such as flight upon encounter, as a reaction to the presence of predators (Kusch 1993; Broglio et al. 2001; Hammill et al. 2009, 2010; Wu et al. 2010). Moreover, resource-dependent morphology has been observed in predatory ciliate species, as they changed their cell volume in response to the size of their prey, and thereby increased the probability of a successful attack (Hewett 1980, 1988). Because of this existing evidence of RAEs and NCEs and the possibility of collecting individual-based morphology, movement and location data through videography, ciliate predator-prey pairs thus offer a convenient and informative system to further study and describe RAEs and NCEs.

We tested for NCEs on prey swimming speed and spatial grouping behavior, hypothesizing that in the presence of the predator the prey would swim faster to increase evasion rates, and would group more and increase in body size to decrease the *per capita* predation risk. Regarding RAEs, we investigated predator body size, swimming speed and diffusion in absence and presence of prey, and the dependence of predator movement on local prey scarcity. We expected the predator to decrease in size due to food scarcity and to swim faster and search in larger areas (as a consequence of swimming faster and straighter) in the absence of prey and to perform bigger displacements the fewer prey individuals are in its proximity to increase the encounter rate with prey.

## Material and Methods

### Material and Experiment

We carried out an experiment to observe the behavior of ciliates *Didinium nasutum* (predator) and *Paramecium caudatum* (prey) in absence and presence of each other. Before and during the experiment, we kept the ciliates in bacterized organic protozoan pellet medium (Carolina Biological Supply Company, Burlington NC; concentration of 0.55 g L^-1^, see Altermatt et al. 2015) at 15 °C. The predator was kept in several maintenance six-well plates and fed with *P. Caudatum ad libitum*.

The experimental units were Petri dishes (diameter of 5.4 cm) each containing 3 mL of bacterized medium and only *P. caudatum* individuals (prey monoculture), only *D. nasutum* individuals (predator monoculture) or both species (mixed culture), with four replicates for each of these three compositions. From the maintenance plates, we randomly selected 60 predator individuals and manually pipetted them to each replicate of the predator monoculture and of the mixed culture. We confirmed the number of predator individuals under a light microscope after the transfer. In the case of the mixed and the prey monoculture we added approximately 400 prey individuals per milliliter. We incubated the replicates for four hours in the dark to give the individuals time to reinstate any behavioral and spatial dependency that might have been disturbed during the preparation of the experimental units.

After the incubation time, we recorded three 30 seconds videos of each replicate at magnification 12.5 with 25 frames per second (750 frames per video) using a Hamamatsu C11440 camera, a Leica M205 C dissecting microscope with dark field illumination, and the software HCImage Live. We had an interval between videos of the same Petri dish of one minute, to ensure a greater independence of the videos as individuals could move in and out of the frame in that time. The videos covered a square of side-length 1.7cm (i.e., an area of 2.89 cm^2^, 12.6% of the surface of the Petri dish) in the center of the Petri dish. The captured area within the Petri dishes was the same across all experimental units and data collection was blinded.

We analyzed the videos with the R-package bemovi (Pennekamp et al. 2015), which tracks particles and extracts their trajectory and morphology data. To ensure data quality, we only analyzed trajectories that were at least two seconds long. We determined species identities of the tracked particles in the mixed culture replicates with a random forest classification (0.04% out-of-bag estimate of the error rate).

### Data Analysis

We used various variables to investigate the research questions and hypotheses. These variables, their calculations and use are described below and summarized in Table 1.

**Table 1.** Variables, their units, how they were calculated and their use (response or explanatory variable)

| Variable | Unit | Calculation | Used as |
| --- | --- | --- | --- |
| **Predator swimming speed** | $\mu\text{m/s}$ | Per trajectory: 60th percentile of recorded local swimming speed maxima within trajectories. Across trajectories: skew normal distribution fit to trajectory swimming speed distribution. | Response variable |
| **Prey swimming speed** | $\mu\text{m/s}$ | Per trajectory: median of recorded swimming speeds within the two-dimensional parts of trajectories. Across trajectories: mixture of two normal distributions fit to trajectory swimming speed distribution. | Response variable |
| **Predator body size** | $\mu\text{m}^2$ | Per trajectory: mean of recorded body sizes. Across trajectories: skew normal distribution fit to trajectory body size distribution. | Response and explanatory variable |
| **Prey body size** | $\mu\text{m}^2$ | Per trajectory: mean of recorded body sizes recoded within the two-dimensional parts of trajectories. Across trajectories: skew normal distribution fit to trajectory body size distribution. | Response and explanatory variable |
| **MSD: Predator displacements** | $\mu\text{m}^2$ | The squared displacements of predators averaged over time (i.e. within trajectories) and over the ensemble (i.e. across trajectories). | Response variable |
| **Local prey and predator abundance** | cells/<br>area | For each individual, the local number of prey and predator was determined as the number of respective individuals within a detection radius of 1500 $\mu\text{m}$. | Response variable |
| **Grouping Index (GI)** | 1 | One minus the empirical mean nearest neighbor divided by the theoretical uniform neighbor distance. The resulting index can have values between 0 (no grouping) and 1 (complete grouping) | Response variable |

#### Swimming speed and body size

The recorded videos were two-dimensional, providing the (*x,y*)-coordinates of the individuals across time. However, while thin (i.e. 1.31 mm) the layer of medium in the Petri dishes still allowed the ciliates to move in the *z*-direction to a small extent. In fact, the predator typically swims in helices (Salt 1979), which is a three-dimensional type of movement. To account for this in the estimation of the predator swimming speed, we made use of fact that the *z*-component of the velocity of a helical trajectory is zero twice per complete helix turn, if the helix lies in the (*x,y*)-plane (see Gurarie et al. 2011). We identified these points as the local maxima of the two-dimensional speed of a predator individual (i.e., its (*x,y*)-displacement per frame). We then estimated the swimming speed of each predator individual by taking the 60th percentile of these local maxima, as this percentile proved to be a reliable estimate of the swimming speed (see supplementary Section S2.1 for more information regarding predator swimming speed calculation). As the predator is approximately the shape of a sphere (Kahl 1930) and thus independent of its orientation, we estimated the body size of each predator individual by taking the mean of its recorded body sizes during its entire trajectory.

Unlike the predator, the prey swims in more or less straight lines. As the prey is cigar-shaped with a length of 170-290 μm and a width of 50-80 μm (based on Wichterman 2012, and personal observations), its aspect ratio is always larger than two when it swims in the (*x,y*)-plane but it can be smaller than two when it swims perpendicularly to the (*x,y*)-plane. We estimated the swimming speed and the body size of each prey individual by respectively taking the median of its swimming speed and the mean of its body size that we recorded considering only the parts of its trajectory in which the aspect ratio was larger than or equal to two (indicating only small or no movement in the *z*-direction, see supplementary Fig. S8).

To test for differences in swimming behavior and body size in absence and presence of their trophic opponent, for both the prey and the predator we selected suitable theoretical distributions (i.e. probability density functions) for their swimming speeds and body sizes based on the empirical distributions of these measures. We then fitted these distributions via maximum likelihood (R-package bbml, Bolker and Team 2017) and compared the estimated parameters and their corresponding 95% confidence intervals between mixed and monocultures. For the swimming speed of the predator as well as for the body size of both species, we chose a skew normal distribution (function snorm() in R-package fGarch, Fernández and Steel 1998; Wuertz et al. 2019). In this distribution, a symmetric normal density is achieved with the skewness parameter = 1, right- and left-skewed normal distributions respectively with skewness > 1 and < 1. For the prey swimming speed, we used a weighted mixture of two normal distributions, i.e. 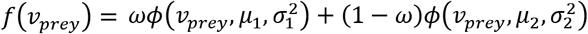. In the equation, *v_prey_* is the swimming speed of the prey and *f*(*v_prey_*) its density function, *ϕ* is the density function of a normal distribution, *ω* the weight used to combine the two normal distributions and *μ*_1_, *μ*_2_, *σ*_1_ and *σ*_2_ respectively the means and the standard deviations of the two normal distributions. To assess the relation between swimming speed (response variable) and body size (explanatory variable) in the two species, we fitted linear models including an interaction with the culture type (mixed and monoculture).

#### Predator displacements

To characterize the displacements of the predator we computed the mean square displacement MSD (i.e., the average squared displacement of the predator) using the estimator 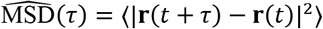 (see e.g. Ariel et al. 2015). In the equation, **r**(*t*) is the location vector at time *t* and the angle brackets indicate the averaging done first over time (time-average) and then over the number of trajectories (ensemble-average) for a time-lag value *t* (see for instance Janczura and Weron 2015). The MSD is further proportional to time (MSD ∝ *t^γ^*, see Méndez et al. 2014) with the exponent *γ* indicating whether displacements follow a normal (*γ* =1) or an anomalous diffusion (*γ* ≠ 1).

To assess whether the predator performed larger displacements in the absence of prey we compared the 95% confidence intervals of the estimated MSD values (based on 10’000 bootstrapped replicates of the calculated time-average MSD). We investigated changes in types of diffusion with linear regression by estimating *γ* as the slope in the log_10_transformed relation between MSD and time for time-lag values in the interval from 0.32 to 9 seconds (from 8 to 225 frames). The lower bound of the interval was chosen such that the helices of the predator trajectories did not influence the MSD, and the upper bound was limited by the number of long trajectories.

We computed the number of neighbors of each predator individual as the number of predators and prey within a detection radius of 1500 μm. We chose this radius considering the swimming speed of the predator and carried out a sensitivity analysis to assess its impact on the analysis. To limit computational costs, we did this only for every 25th recorded video frame. The results of the analysis were invariant to which frames were selected. We further computed the displacements that the predator individuals made in between the considered frames.

We investigated the relation between predator displacements (response variable) and prey abundances with a linear regression. The independent variables were prey abundance, predator abundance, and the culture type (i.e., predator monoculture or mixed culture) with an interaction term between the latter two. Further, to assess whether the number of neighbors (regardless of species identity) or the prey absence had a bigger impact on predator behavior, we calculated the mean number of neighbors for the predator individuals tracked in the local neighbor analysis described above. We then used this as an independent variable together with an interaction with culture type in a linear regression that had the swimming speed of the considered predator individuals as the response variable.

#### Prey grouping

We computed a grouping index to quantify the effect of the predator on the degree of grouping in the prey. For every 25th frame we counted the number of observed prey individuals *n* and calculated the Euclidian nearest prey neighbor distance of each prey individual present. We then averaged the nearest prey neighbor distance within each frame to yield the mean nearest neighbor distance, denoted here as MNND.

For comparison, we estimated the uniform neighbor distance UND(*n*) (i.e. the distance between *n* equally spaced individuals, see supplementary Fig. S13). We did this estimation by using the function rSSI() in the R-package spatstat, which places *n* individuals in the area captured by the videos according to a given inhibition distance (the individuals cannot be placed closer to each other than this distance). We selected the biggest inhibition distance as the UND(*n*) for which the function succeeded in placing *n* points. To ensure that we found the biggest possible UND(*n*), we called the function up to 2500 times or until the *n* points were successfully placed for each inhibition distance and we terminated each call after 1000 failed point placements.

We defined the grouping index GI as GI(n) = 1 - MNND/UND(*n*) and calculated it for each considered frame. By dividing MNND by UND(*n*), the index is standardized for the different number of individuals *n* present in the frames. The resulting index can have values between 0 (no grouping, the individuals have an inhibition distance) and 1 (complete grouping, all individuals are in one point) and accounts for different number of individuals present, which means that GI is comparable across frames.

We used the grouping index GI as the response variable in a normal mixed effects model with the culture type (mixed and monoculture), time and their interaction as fixed effects and video nested in sample as random intercepts. As a test of robustness, we repeated this analysis with a different measure of prey grouping as the response variable (based on the intensity functions of prey spatial point patterns, see supplementary Section S4). Further, the analysis was robust to the selection of the frames considered.

## Results

We tracked a total of 11423 trajectories that were at least two seconds long. Of these, 14.7% belonged to predator individuals and 85.3% to prey individuals. Most trajectories were shorter than 10 seconds (Fig. S1). After the incubation period of four hours, there were fewer prey cells in the mixed culture replicates than in the prey monoculture replicates, suggesting either predation of *P. caudatum* by *D. nasutum* or decreased growth rate of the prey in presence of the predator, or both (Fig. S2).

### Predator swimming speed and body size

In both the monoculture and the mixed culture, the predator revealed two distinct swimming regimes (supplementary Fig. S6a). This was further confirmed by a hierarchical cluster analysis of the trajectories (see Section S2.1). Hence, we classified predator individuals as either slow or fast, depending on whether it’s swimming speed was less or greater than 1500 μm s^-1^, respectively.

We fitted the skew normal distributions separately to the swimming speed data of the fast and the slow predator (Figs. 1a,b). On average, slow-swimming and fast-swimming predator individuals swam respectively 6.3% and 16.6% slower in the mixed culture than in the monoculture (non-overlapping 95% confidence intervals, see Fig. 1c and Table S3). This difference was independent from the length of the trajectories considered (Fig. S7). There was significantly less variation in swimming speed for the fast predator in monoculture when compared to the mixed culture, while no other differences in the standard deviation and the skewness of the swimming speed distributions where present (Figs. S6e,f).

**Fig. 1.**
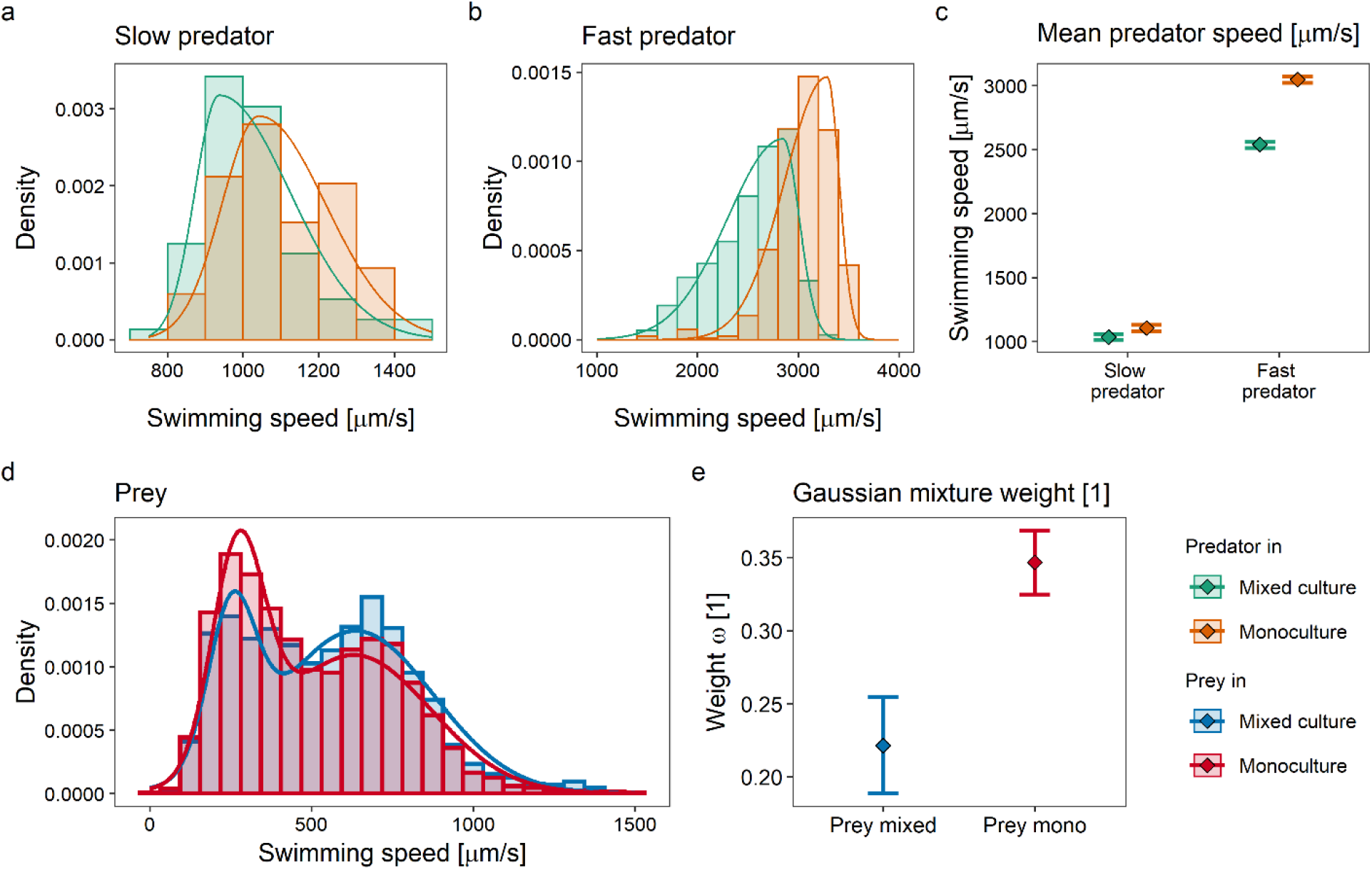
**a – b** Empirical and fitted skew normal densities for the swimming speed of the predator in the two different cultures, respectively for the slow (**a**) and the fast (**b**) individuals. **c** Estimated mean swimming speed of the slow and the fast predator in mixed and monoculture. **d** Empirical and fitted Gaussian mixture densities for the swimming speed of the prey in the two different cultures. **e** Estimated Gaussian mixture weights for the swimming speed distribution of the prey in mixed and monoculture. In **c** and **e** the error bars indicate 95% confidence intervals

The body size of the fast-swimming predator significantly increased by 17.7% in the mixed culture (non-overlapping 95% confidence intervals, see Figs. 2b,f and Table S9). This was not the case for the slow predator, where the increase by 4.1% in body size in the mixed culture was not significantly different from 0 (overlapping 95% confidence intervals, see Figs. 2a,f). Further, the swimming speed of the slow predator increased by 43.7 μm s^-1^ (*t*-value=7.04, df=266, *p*-value<0.001) for each 1000 μm^2^ increase of its body size, with no difference between culture types (*t*-value=0.87, df=266, *p*-value=0.385, see Fig. 2d and Table S10). There was no significant relation between body size and swimming speed (*t*-value=0.45, df=1400, *p*-value=0.652) for the fast predator, while it was confirmed that it swam slower in mixed culture (*t*-value=5.73, df=1400, *p*-value<0.001).

**Fig. 2.**
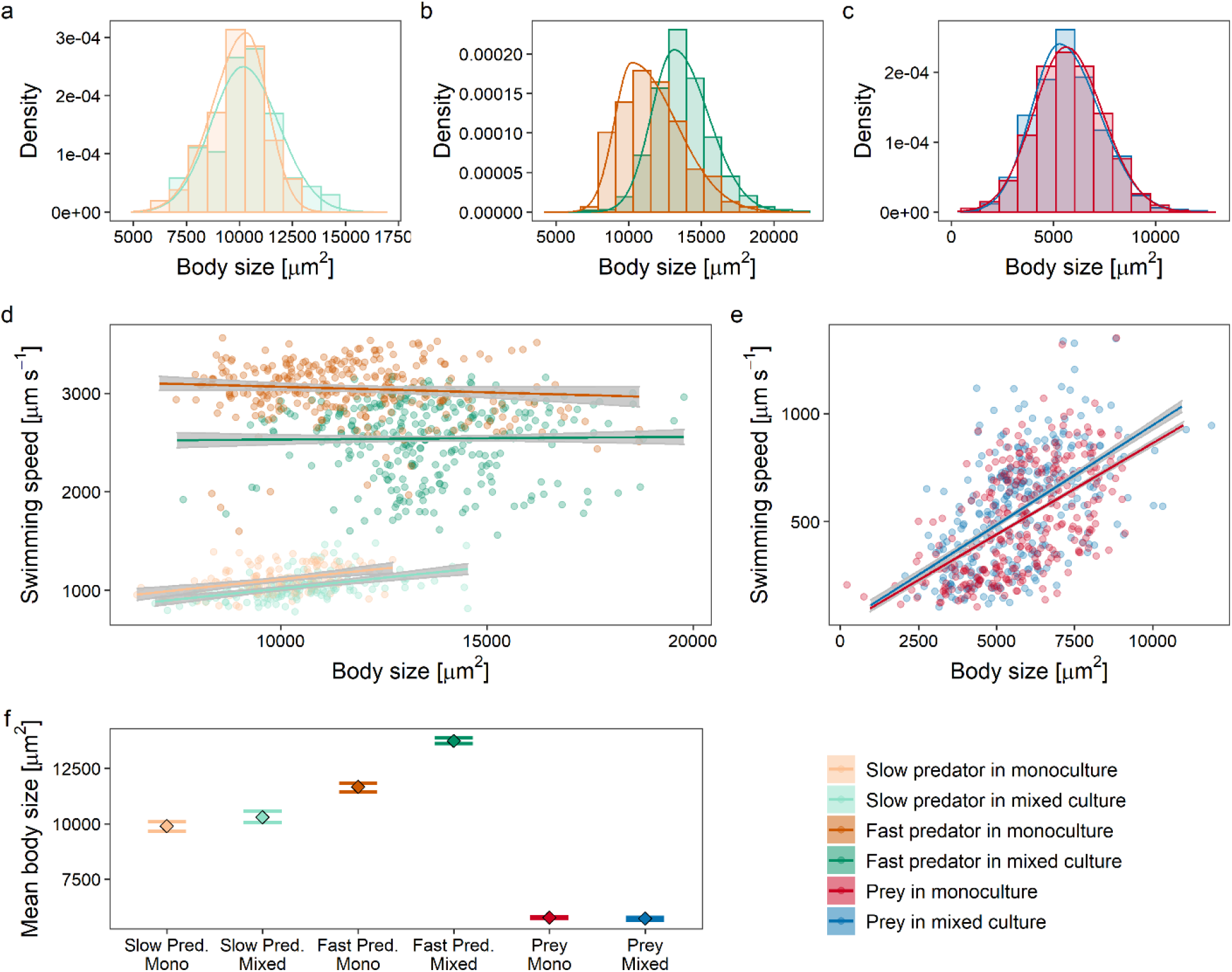
**a – c** Empirical and fitted skew normal densities for the body size of the two species in the two different cultures, respectively for the slow predator (**a**), the fast predator (**b**) and the prey (**c**). **d – e** The relation between species body size and swimming speed in the two types of culture for the slow and the fast predator (**d**) and the prey (**e**). To avoid over-plotting, we randomly selected 300 data points of each group to be displayed in a scatter-plot. Solid lines indicate regression fits and the shaded areas represent the pointwise 95% confidence intervals. **f** Estimated mean body sizes of the prey and the predator in presence and absence of each other. The error bars indicate 95% confidence intervals

### Prey swimming speed and body size

The prey showed two swimming speed regimes also, as the distribution was bimodal (Fig. 1d). The proportion of prey individuals showing the fast movement behavior increased by 12.5% from 65.3% to 77.8% in the presence of the predator (non-overlapping 95% confidence intervals in Fig. 1e and Table S4). We found this difference also for trajectories of only a certain length range (Fig. S10) and also when for the prey mixed culture we only considered videos in which the prey abundance was comparable to the prey abundances in the monoculture (Fig. S2 and Fig. S11). The mean slow swimming speed (i.e., *μ*_1_) was 7.8% greater in the monoculture compared to the mixed culture, the other parameters were not significantly different between the two cultures based on their 95% confidence intervals, (Figs. S9b,c).

The prey body size was not significantly affected by the presence of the predator (overlapping 95% confidence intervals, see Figs. 2c,f and Table S9). For every increase in body size of 1000 μm^2^ the prey swam 84.3 and 92.2 μm s^-1^ faster (*t*-value=36.46, df=9707, *p*-value<0.001). respectively in mono- and the mixed culture, with a significant difference between the two slopes (*t*-value=-2.64, df=9707, *p*-value=0.008, see Fig. 2e and Table S10).

### Predator displacements

The predator displacements were smaller in the presence of prey (non-overlapping confidence intervals in Fig. 3a) and showed superdiffusion in all cases (slope *γ* > 1 in Fig. 3b). The MSD values after one second were 27.0% and 12.5% smaller in mixed culture, respectively for the slow- and the fast-swimming predator (Table S5). After 9 seconds (the largest time-lag value considered for the MSD) these values were respectively 35.9% and 8.4% but with overlapping 95% confidence intervals in the case of the fast-swimming predator likely caused by the small number of trajectories that were 9 seconds long or longer.

**Fig. 3.**
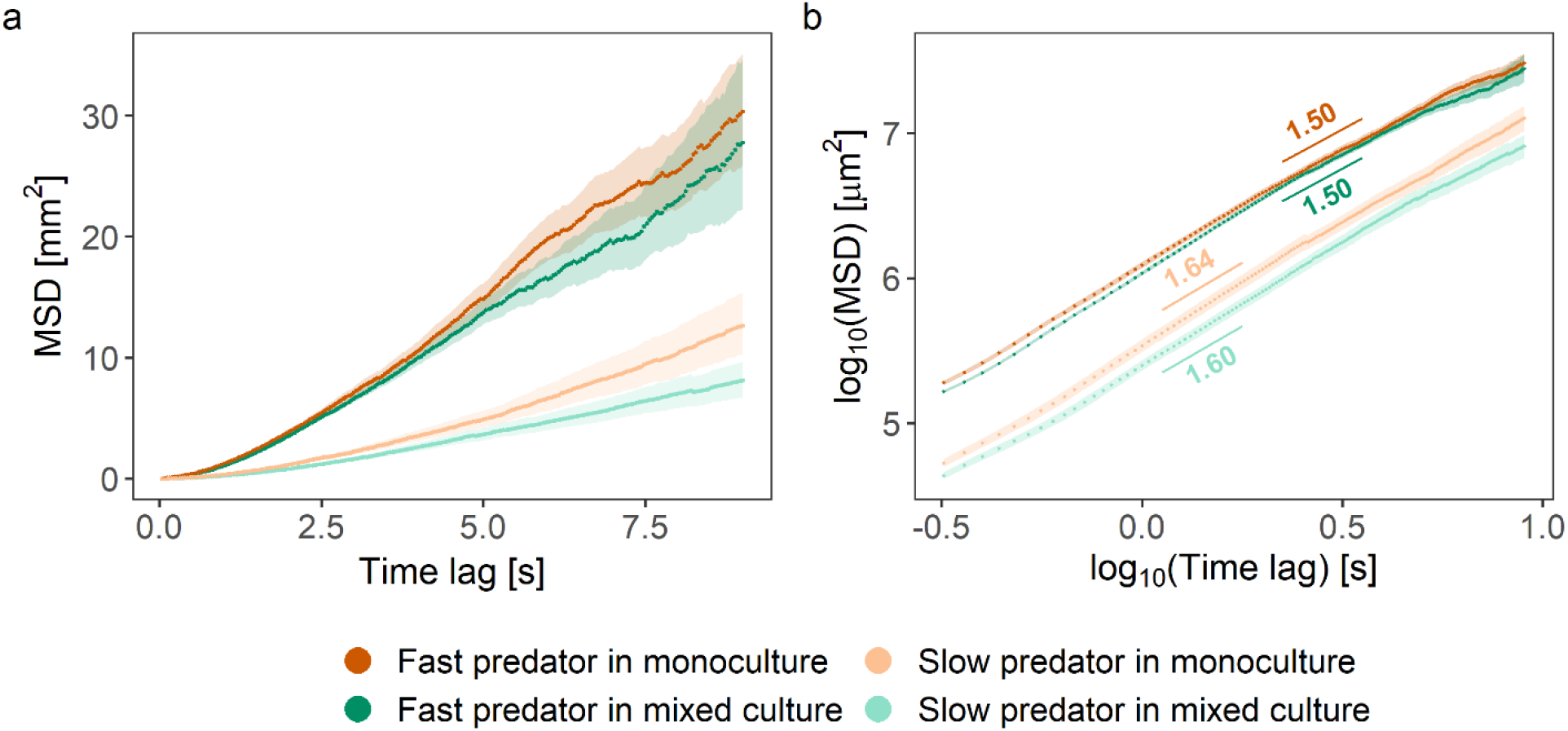
Mean squared displacement (MSD) of the predator as a function of time, for both the predator monoculture and the mixed culture, as well as for both the fast and the slow predator. Bootstrapped pointwise 95% confidence intervals are included (shaded areas). **a** On normal axes. **b** On log10-transformed axes with corresponding slope estimates displayed

Predator individuals also showed displacements that were dependent on the local number of neighbors (Fig. 4). Both the slow- and the fast-swimming predator made smaller displacements the more local prey neighbors they had, with the latter also making smaller movements when it had more predator neighbors in the monoculture (Fig. 4b). Quantitatively, with respect to its biggest displacement the predator made 2.3% (*t*-value=-2.70, df=8946, *p*-value=0.007) and 3.7% (*t*-value=-3.28, df=3918, *p*-value=0.001) smaller displacements for each unit increase in number of local prey neighbors, respectively for the fast- and the slow-swimming predator (Table S6). These patterns persisted in the sensitivity analysis of different neighbor detection ranges (Table S6). The analysis of predator swimming speed showed that the predator swam faster in the monoculture than in the mixed culture regardless of number of neighbors, suggesting that the presence or absence of prey had a larger impact on predator swimming speed than the number of neighbors (see supplementary Fig. S12).

**Fig. 4.**
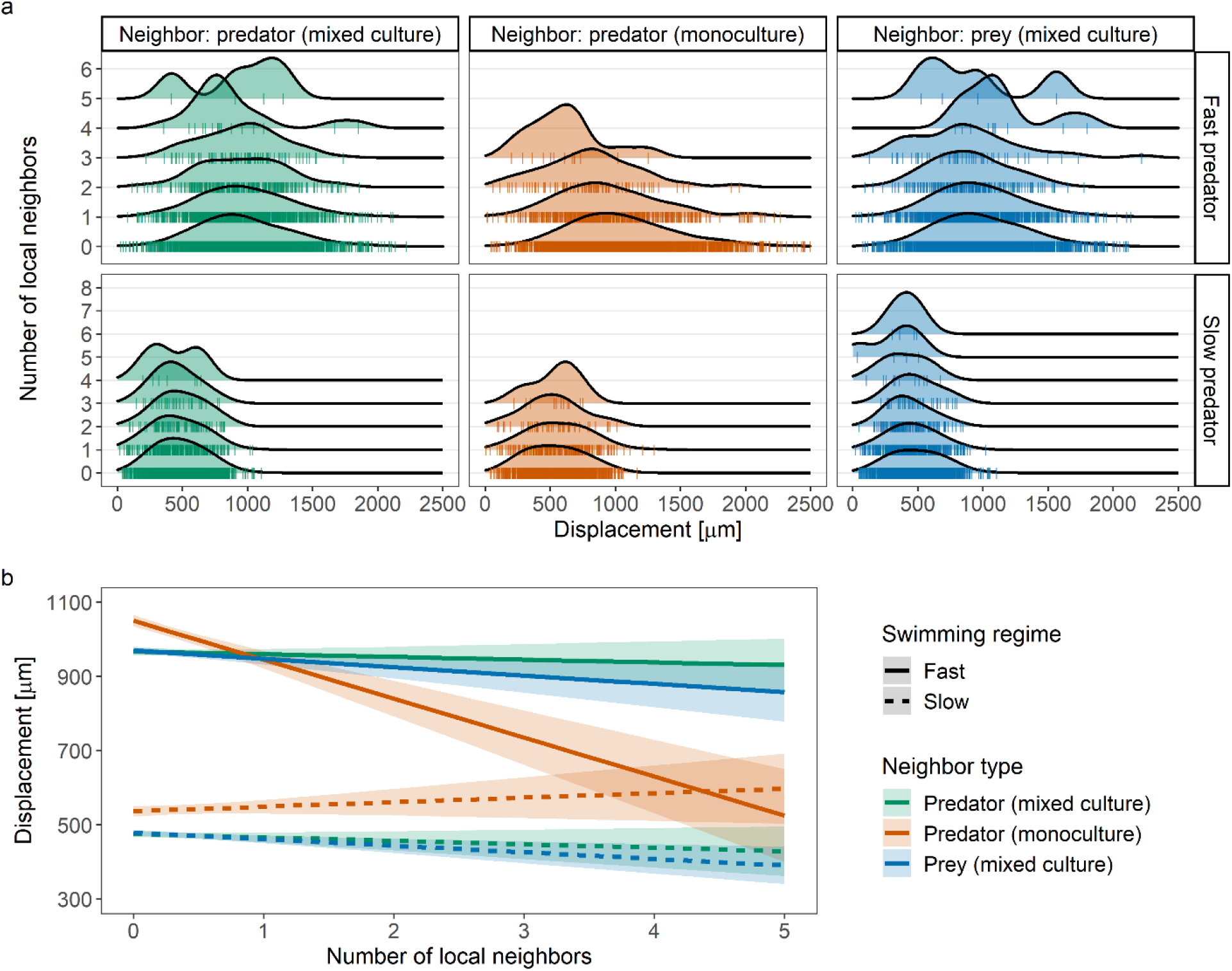
Predator displacement per second as a function of the local number of neighbors at the start of the displacement. **a** Ridge-line plots of the data including rug plots. The top row shows the fast-swimming predator, the bottom row the slow-swimming predator. The columns display the data for when the neighbors were predator individuals (columns one and two, respectively for the mixed and the monocultures) and prey individuals (third column). **b** Results of the linear regression (including 95% pointwise confidence intervals). The number of prey neighbors was set to zero if the effect of predator neighbors was considered and *vice versa*. Solid lines represent the fast-swimming predator, dashed lines the slow-swimming predator

### Prey grouping

On average the grouping index was larger in the mixed culture replicates, but it also showed considerable variability in time (Fig. 5a). The mixed effects model revealed evidence for increased grouping in presence of the predator, with the prey grouping 7.8% more in the mixed culture at time 0 (*t*-value=-4.54, df=713, *p*-value<0.001, see Fig. 5b and Table S8). However, the 95% confidence intervals overlapped in later parts of the videos, suggesting that the relation between predator presence and prey grouping cannot be clearly answered with this data. The alternative analysis based on intensity functions yielded comparable results (see Fig. S15 and Table S8). Further, no difference in prey grouping was found when only videos were considered in which the prey abundance was comparable between the mixed and the monoculture (Fig. S16).

**Fig. 5.**
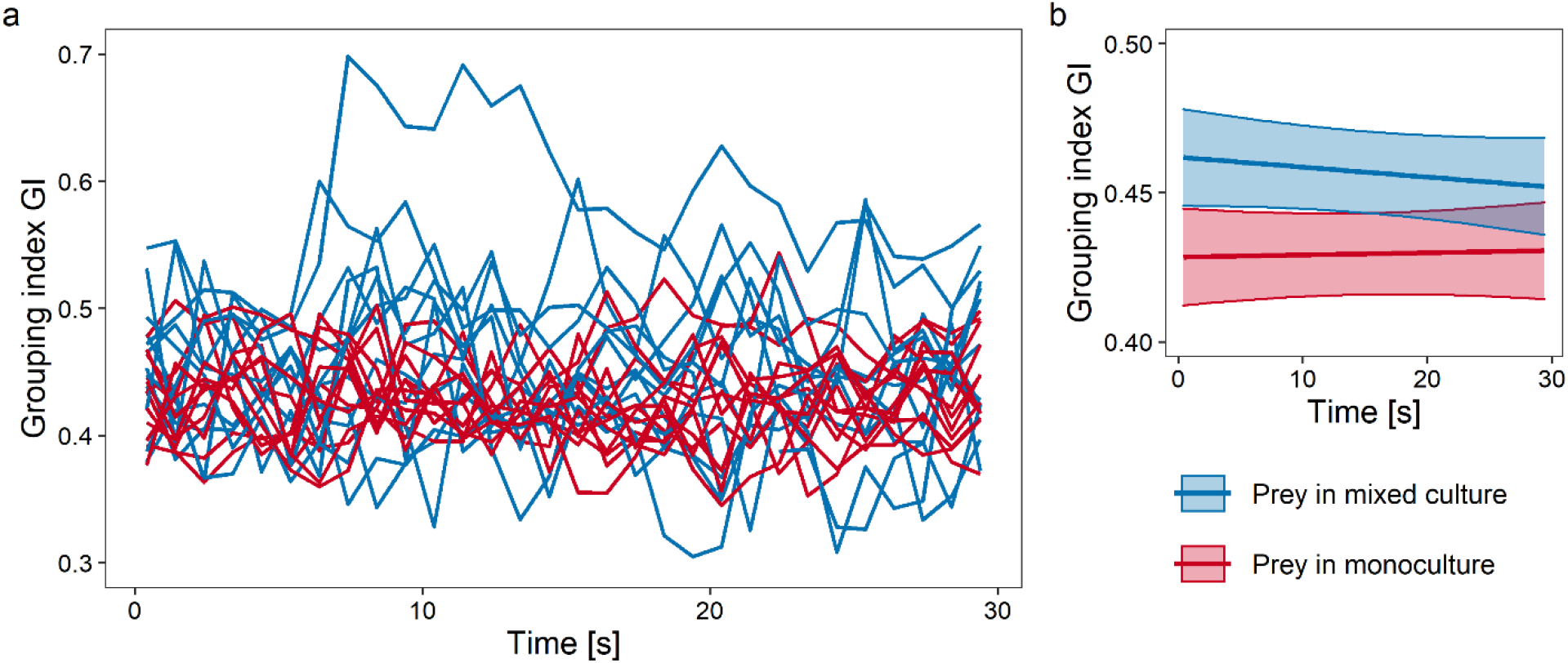
Prey grouping factor as a function of culture type and time. **a** The calculated data, grouped by videos (lines). **b** Results of the mixed effects models fitted. The pointwise 95% confidence intervals are based on fixed effects only (shaded area)

## Discussion

We found evidence of both non-consumptive effects (NCEs) of predators on prey, and of resource availability effects (RAEs) of prey on predators. One NCE was the proportionally greater number of prey individuals swimming faster in the presence of predators. RAEs included the presence of prey causing predators to significantly decrease swimming speed and displacements, and to increase body size. Predator displacements were also dependent on local prey density. Together, these results suggest that the predator counteracts the prey scarcity by changing its behavior to search more quickly and in larger areas, in a likely attempt to increase the encounter rate with the prey, while the prey reacts to high predator densities by swimming faster and thus by possibly increasing the evasion rate upon encounter with the predator, though while also likely increasing encounter rates.

NCEs are a frequently studied component of predator-prey interactions (see for instance Lima 1998; Werner and Peacor 2003; Hermann and Landis 2017; MacLeod et al. 2018). The change in behavior of the average prey individual being more likely to exhibit the fast-swimming speed in the presence of the predator is confirmed by similar results (e.g. Broglio et al. 2001; Hammill et al. 2009) and represents evidence for flight behavior, as it was presumably made by the prey to increase its escape ability upon encounter with the predator (Sih 2011). Flight as an anti-predator behavior is a widespread phenomena (see for instance Krause and Godin 1996; Stankowich 2009; Harvey and Menden-Deuer 2012) and has also been reported for ciliates (Tamar 1979). As the predator was relatively abundant in the mixed cultures, it is further possible that the prey was in a constant state of flight behavior. This could potentially also explain the only weak evidence for increased prey grouping caused by predator presence, a behavior often observed in animals (e.g. tadpoles and cercopithecoid primates, see respectively Hill and Lee 1998; Spieler 2003).

Contrasting this, a lot less is known about RAEs. The investigation of trait mediated interactions in predator-prey systems is most often centered around the prey and how it evades or escapes the predator (see for example Wahl and Stein 1988; Miller et al. 2014) and much less frequently on the predator and how it changes its search strategy at low resource availability (e.g. Munk 1995; Ronconi and Burger 2008). Rare evidence for an increased predator swimming speed in the absence of prey was reported by Kuefler et al. (2013), who similar to our results found that on average their studied rotifer species swam faster in absence of food but only when there were no conspecific individuals present. Assuming that increased movement rates and displacement lengths are likely to be associated with a better chance of encountering prey, the here reported increase in swimming speed and displacements represents evidence that the predator *Didinium nasutum* is capable of responding to increased hunger levels by not only covering more space but also by searching in it more quickly. On this note, larger predator displacements as a counter to the lack of food, either to search in larger areas or to move to other patches altogether, have been reported in the context of Lévy flights (a particular search strategy, see e.g. Humphries et al. 2010; Sims et al. 2012).

With the presence of both RAEs and NCEs confirmed in such a widely studied predator-prey pair, it becomes evident that neglecting these effects when the interaction strength between species is assessed is questionable. However, when the interaction strength is simplified to correspond to the functional response, such neglect is caused, i.e. it is assumed that prey intrinsic growth rate is independent from the predator density and that predator conversion efficiency and mortality rate are invariant to prey availability (Rall et al. 2010; Sentis et al. 2012; Kalinoski and DeLong 2016; Daugaard et al. 2019). Yet, it has long been known that prey growth does, in fact, depend on predator density (Preisser et al. 2005; Clinchy et al. 2013) while it has recently also been pointed out that the conversion efficiency and the mortality rate are associated with prey abundance (Minter et al. 2011; Li and Montagnes 2015). Crucially, the functional response approach only estimates the consumptive effects of predators on prey and can only partially and implicitly incorporate NCEs and RAEs. For example, while induced defensive strategies may cause a decrease in the predator feeding rates and are thus at least in part captured by the functional response parameters, this approach does neither incorporate the growth rate of the prey nor any (e.g. predator-mediated) changes in it. Ultimately, concrete knowledge about predation effects other than predator feeding rates, such as changes in prey behavior in predator presence, has the potential of providing insights into the mechanisms that influence the functional response characteristics of the system (e.g. whether the predator exerts higher or lower predation pressure on the prey when the prey is only present in small numbers).

The estimated predator swimming speed revealed two clearly different movement rate regimes (i.e., the slow- and the fast-swimming predator). The only other study finding similar results did so in a heavily altered medium (in which the average swimming speed reached only 3% of the swimming speed in the unaltered medium, see Luckinbill 1973). It is possible that the more modern videography approach used in our study was the factor that allowed the detection of the two swimming speed regimes in a more natural setting that has been overlooked so far and that represents a gap in the understanding of *D. nasutum*.

A potential explanation for the presence of this bimodal swimming behavior could be that individuals swimming slower are the ones that recently went through reproduction, which would explain why there are proportionally many more fast individuals than slow ones (Table S1). Considering that the slow predator was generally also smaller than the fast predator in both the mixed and the monocultures (Fig. 2f), this further suggests that the slow predator recently went through cell division (i.e. reproduction). More evidence in favor of this is that only the body size of the fast predator decreased in the absence of prey (i.e. due to starvation), while the slow predator did not significantly change in size — it is possible that only individuals of a certain size go through cell division. The analysis of the relation between morphology and speed also revealed that the swimming speed of both prey and slow predators increased with body size (Figs. 2d,e). These results suggest that increasing the body size leads to higher swimming speeds, but with diminishing returns as the movement of the fast predator was independent from body size.

While the described NCEs and RAEs for this species pair are likely to influence the prey-predator interaction strength and thus the population dynamics of the two species, we do not quantify this here. This represents an interesting research question for further studies, that could for instance extend the work of Griffiths et al. (2018) in which a trait-demography feedback with predator-dependent prey body size was modeled.

Jointly considered, the results show that both the predator *Didinium nasutum* and the prey *Paramecium caudatum* are able to behaviorally and morphologically (only *Didinium nasutum*) react to changes in abundance of their trophic opponent. This is expected, as to maximize their fitness organisms must find a balance between foraging and avoiding being eaten by a predator and therefore the behavior of both the prey and the predator depends on the environment and on its heterogeneity (which includes other species, see Sih 2011). These changes in behavior are likely to cause changes in energy expenditure (Preisser et al. 2005; Clinchy et al. 2013), either to increase predation success or to boost the chances of predation evasion. There is evidence that NCEs do affect the stability of the communities the predator-prey pair is a part of (Anholt and Werner 1995; van Veen et al. 2005; Peckarsky et al. 2008; Fill et al. 2012), but it remains to be determined what role RAEs play in this regard and thus also how big the joint impact of consumptive effects, NCEs and RAEs is on community and ecosystem stability.

## Acknowledgments

Reinhard Furrer and Owen L. Petchey had equally significant roles in the project, so are to be considered joint senior authors and their relative position in the list of authors should be disregarded.

## Author contributions

UD, RF and OLP conceived the ideas and designed the methodology. UD collected and analyzed the data and led the writing of the manuscript. All authors contributed critically to the drafts and gave final approval for publication.

## Ethics declarations

### Ethical approval

This is a study with protists and as such no ethical approval is required.

## Data availability

Data available from the Zenodo repository: https://doi.org/10.5281/zenodo.5676309 (Daugaard et al. 2021)

