## Supplementary_information for "Prey speed up, predators slow down: non-consumptive effects on movement behavior of a ciliate predator-prey pair"

#### Keywords

### Supplementary material

#### S1 Ciliate abundances and trajectories

A total of 11423 trajectories that were at least two seconds long were tracked. Out of these, 14.7% belonged to predator individuals and 85.3% to prey individuals. The distributions of the predator and prey trajectory lengths are shown in Fig. S1. The trajectories were rarely 750 frames (i.e., 30 seconds) long and were very often shorter than 250 frames (i.e., ten seconds).

The number of fast and slow swimming predator trajectories in the mixed and in the monoculture are reported in Table S1. The proportion of fast predators was slightly bigger in the mixed culture when compared to the monoculture (85.4% vs. 81.36%, $\chi^2 = 4.46$, $p$-value $= 0.035$).

Based on observed counts, the abundances of predator individuals after the incubation time was fairly similar in the mono- and the mixed culture (Fig. S2). This was not the case for the prey, were the abundances were clearly lower in the mixed culture, with the exception of two videos (labelled in Fig. S2).

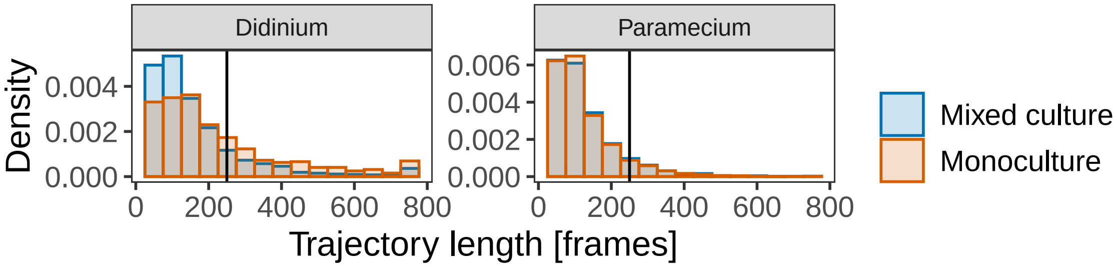


**Fig. S1** Histograms of the trajectory lengths for both the predator and the prey in the mixed culture and their respective monoculture. Vertical black lines denote the ten seconds mark

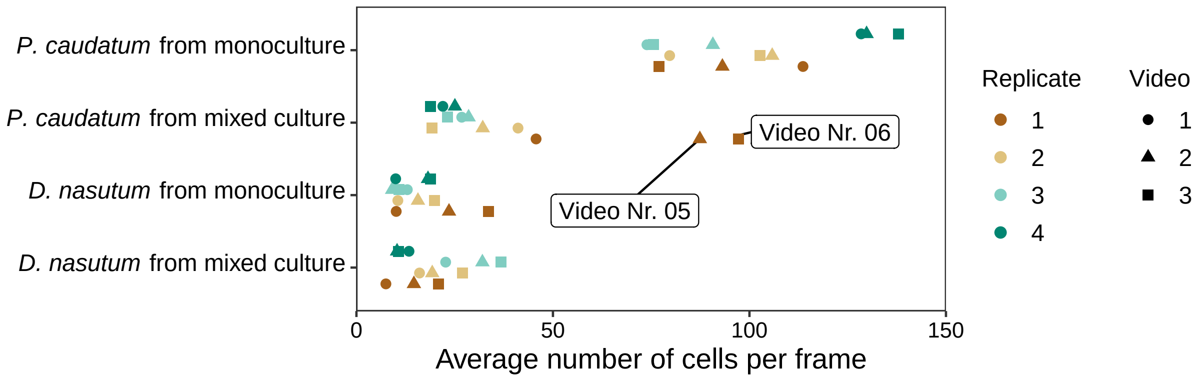


**Fig. S2** Average predator and prey counts per frame in the different replicates and cultures. Points of the same color indicate values from different videos from the same replicate. Within a replicate the videos are shown with different shapes. In two videos of the mixed cultures the prey counts where more comparable to the counts in the monoculture. These two videos are labelled in the figure

**Table S1** Counts of tracked predator trajectories in the mixed and the monocultures both for the fast and for the slow predator

| | Fast predator | Slow predator |
|---|---|---|
| Mixed culture | 889 | 152 |
| Monoculture | 515 | 118 |

#### S2 Ciliate swimming speed and body size

##### S2.1 Predator swimming speed

The velocity vector $\boldsymbol{V}$ of an object is given as $\boldsymbol{V} = \lim_{\Delta t \to 0}(\boldsymbol{r}(t+\Delta t) - \boldsymbol{r}(t))/\Delta t$, with $\boldsymbol{r}(t)$ being the location vector at time $t$, and $\Delta t$ a time amount tending to 0. As the videos were recorded with 25 frames per second, in this study the smallest difference in time was $\Delta t = 0.04$ seconds. The swimming speed $v$ of an individual is then given as the magnitude of the velocity, i.e., $v = |\boldsymbol{V}| = (V_x^2 + V_y^2 + V_z^2)^{1/2}$.

The videos recorded were two-dimensional, providing the $(x, y)$-coordinates of the individuals. However, while thin (i.e. $1.31\,\mathrm{mm}$) the layer of medium in the Petri dishes still allowed the ciliates to move in the $z$-direction to a small extent. Therefore we made the following calculations for predator and prey swimming speed.

The predator typically swims in helices. Figs. S3a,b show a two-dimensional representation of the trajectories of two predator individuals (one fast- and one slow-swimming). The time series of their swimming speed based on the two-dimensional trajectories show a clear periodicity (Figs. S3c,d), with the local maximas denoting where the swimming speed in the third dimension was zero or close to being zero.

**Table S2** Three swimming speed estimates of the predator in the two identified cluster-groups: based on the median of all estimated swimming speeds (ignoring the $z$-component of the velocity), based on the median of the local maxima of the swimming speeds and based on the 60th percentile of the local maxima of the swimming speeds

| Cluster group | Median 2D speed | Median speed | 60th percentile speed |
|---:|---:|---:|---:|
| 1 | 818.52 | 1011.82 | 1067.37 |
| 2 | 1986.76 | 2598.87 | 2763.54 |

Of the predator trajectories, 250 were randomly selected and then trimmed to just contain their first 100 frames. These time series were then used in a hierarchical cluster analysis using Dynamic Time Warping (DTW, Berndt and Clifford 1994) as distance measure and Ward's agglomeration method as the dissimilarity measure. This analysis revealed two distinct groups (Fig. S4 shows the dendrogram of the analysis for when only 50 trajectories were used) that differed in their swimming speed (Table S2), confirming the presence of two swimming speed regimes in the predator.

The $z$-component of the velocity ($V_z$) of a helical trajectory is zero twice per complete helix turn, if the helix lies in the $(x, y)$-plane (Gurarie et al. 2011). These points of zero z-velocity were identified as the local maxima of the $(x, y)$-speed $(V_x^2 + V_y^2)^{1/2}$ across time. For each predator trajectory a single value for its swimming speed was calculated. This was done by taking the 60th percentile of the local maxima of the swimming speed time series (that were calculated based on the two-dimensional data and thus ignore the third dimension). Fig. S5 shows the first 100 frames of 30 randomly chosen predator swimming speed time series. As can be seen, this percentile captured the predator swimming speed for both the fast and the slow predator.

The swimming speed distributions of the predator are shown in Fig. S6. The estimated parameters of the skew normal fits to the predator swimming speed distributions are listed in Table S3, including their 95% confidence intervals. Notably, the mean of the swimming speed was estimated to be significantly bigger in the monoculture than in the mixed culture for both the fast and the slow predator. The predator swimming speed was larger in the monoculture regardless of trajectory length, as indicated by the local polynomial regression shown in Fig. S7.

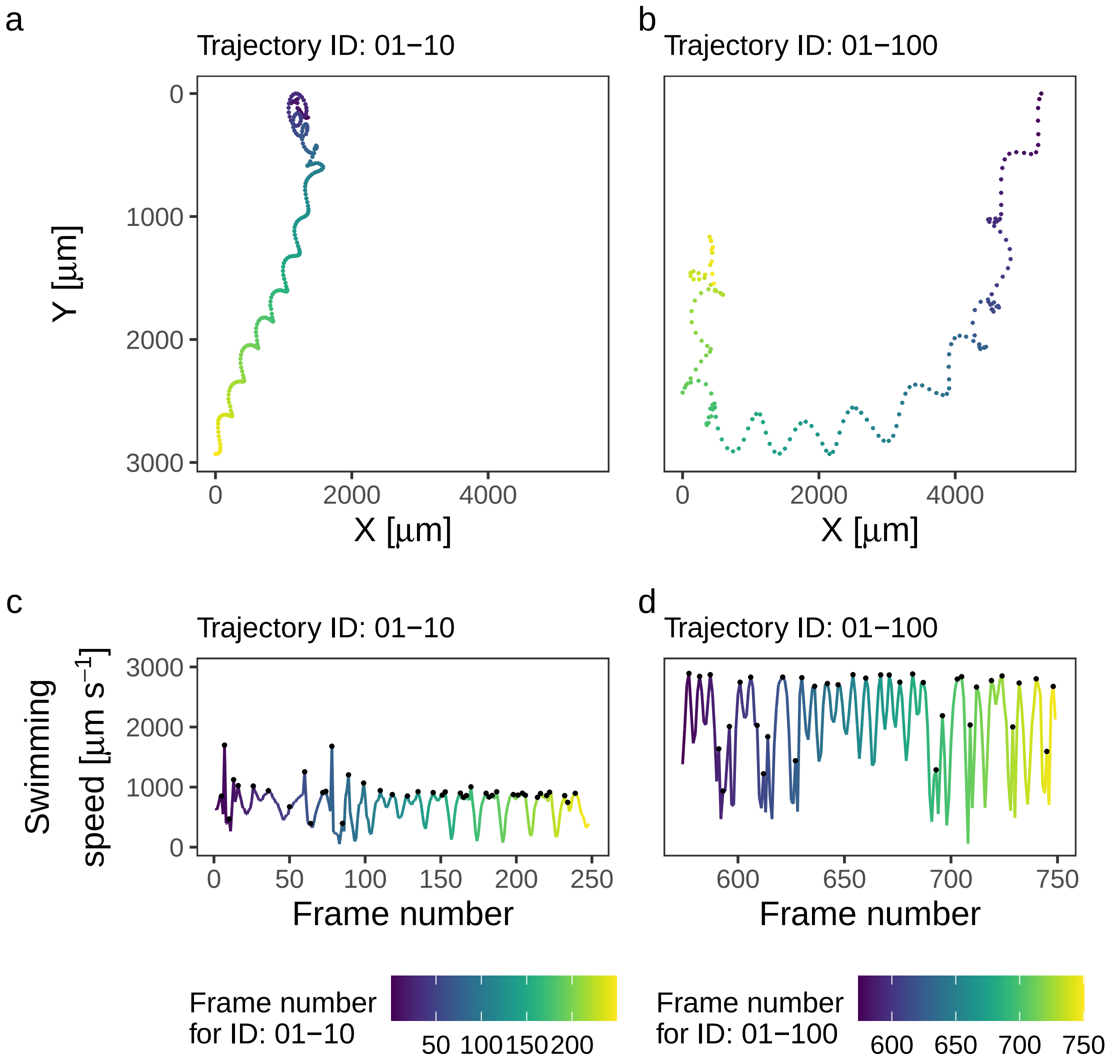


**Fig. S3** Two examples of predator trajectories with IDs 01-10 (**a**) and 01-100 (**b**) and their line graph with frame number on the $x$-axis and swimming speed (assuming $V_z = 0$) on y-axis (respectively **c** and **d**). The trajectories are shifted so that the axes are the same in both trajectory plots. The black dots indicate the local maxima of the swimming speed. The lines are color-coded based on the frame number, i.e., based on the trajectory length

##### S2.2 Prey swimming speed

The distribution of the aspect ratios of prey individuals in the different cultures is shown in Fig. S8a, while Fig. S8b shows exemplary the trajectories captured in a video of a prey monoculture. The individual parts of the trajectories are colored based on whether the aspect ratio was below two or not at that time point. As can be seen, the aspect ratio remained equal to or above two for most of the time, hence only a small part of the trajectories were filtered out

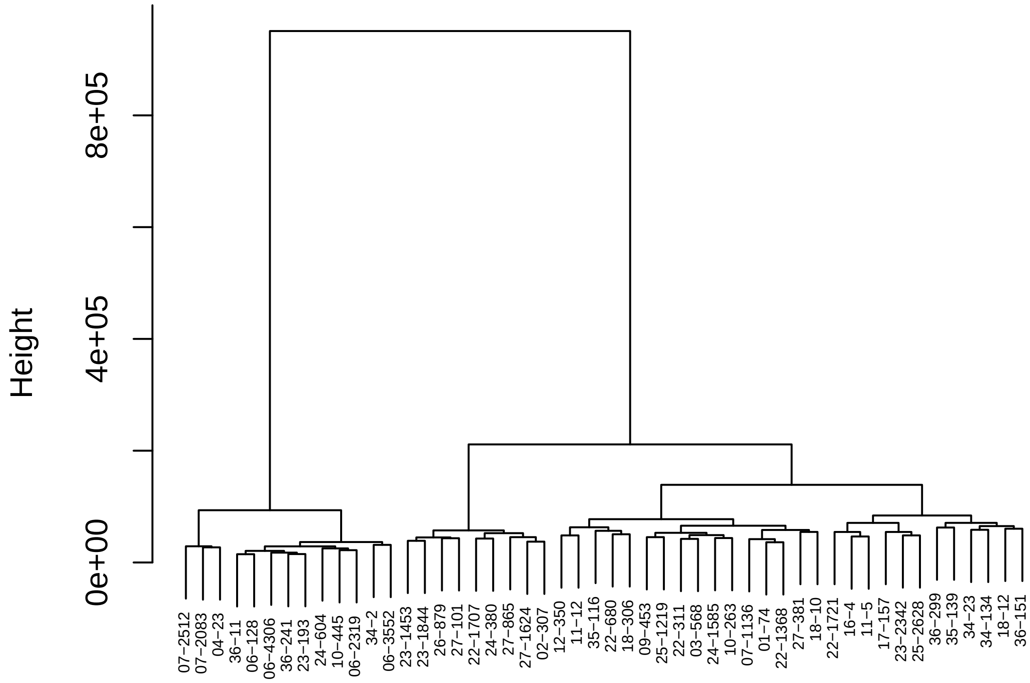


**Fig. S4** Time series clustering of 50 randomly selected predator trajectories. The labels are the IDs of the trajectories

**Table S3** Maximum likelihood estimates of the mean, the standard deviation and the skewness of the swimming speed distribution of the predator based on a skew normal distribution. The estimates are given for both the slow and the fast moving predator in both the mixed and the monoculture

| Culture | Movement | Parameter | Estimate | 95% CI |
|---|---|---|---|---|
| Mono | Fast | Mean | 3046.742 | (3021.75, 3071.57) |
| | | Standard deviation | 289.117 | (269.76, 308.58) |
| | | Skewness | 0.535 | (0.45, 0.64) |
| | Slow | Mean | 1104.438 | (1078.14, 1129.47) |
| | | Standard deviation | 139.722 | (123.07, 162.25) |
| | | Skewness | 1.337 | (0.93, 2.00) |
| Mixed | Fast | Mean | 2539.503 | (2511.78, 2560.37) |
| | | Standard deviation | 377.548 | (360.58, 397.90) |
| | | Skewness | 0.541 | (0.49, 0.60) |
| | Slow | Mean | 1034.650 | (1014.31, 1056.99) |
| | | Standard deviation | 132.184 | (118.09, 149.84) |
| | | Skewness | 1.706 | (1.35, 2.30) |

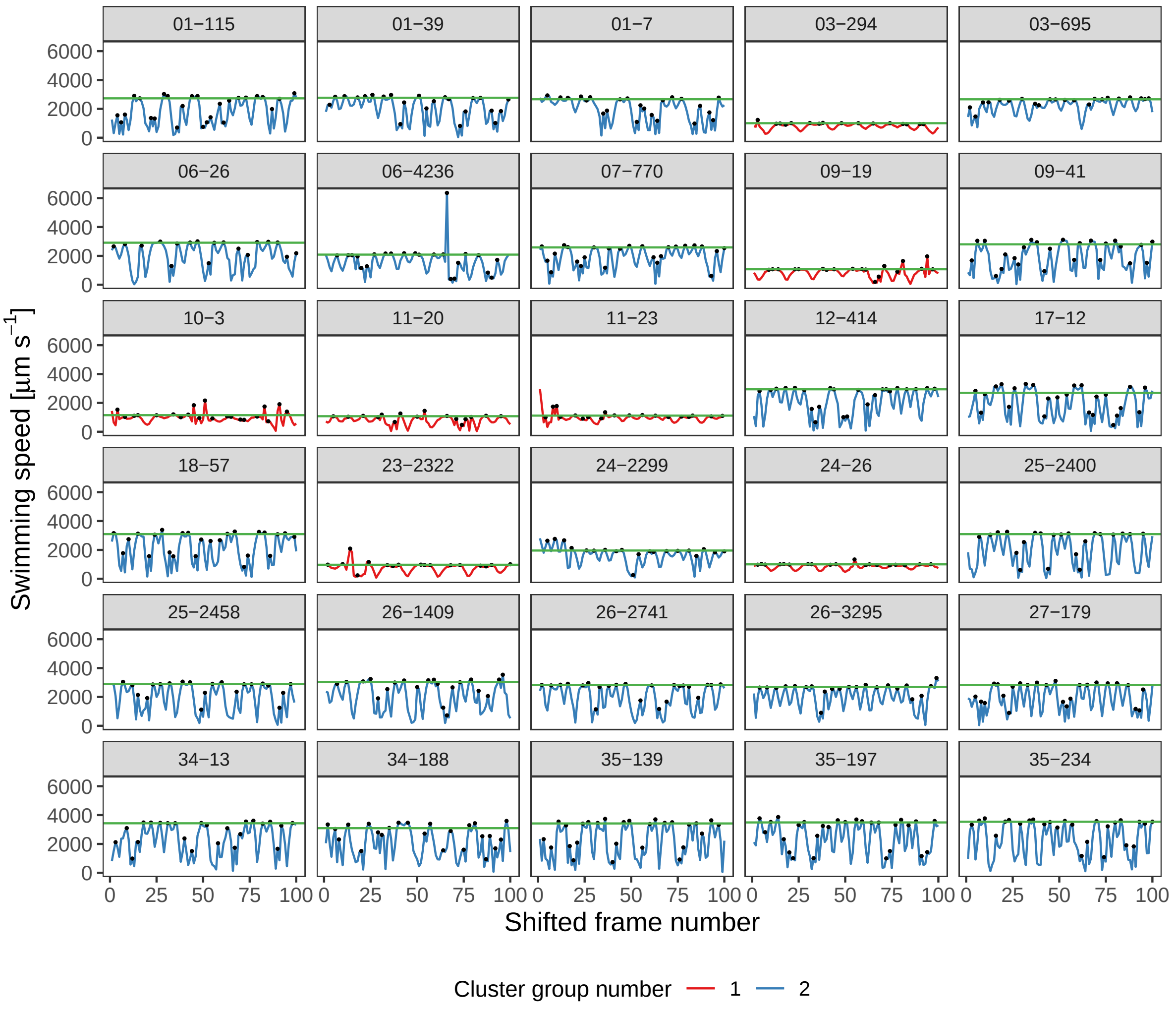


**Fig. S5** The first 100 frames of 30 randomly chosen trajectories, colored according to their cluster group number. The black dots indicate local maxima, and the horizontal green lines indicate the 60th percentile of the local maxima of each trajectory

when the prey swimming speed was calculated. The swimming speed distributions of the prey are shown in Fig. S9. The estimated parameters of the Gaussian mixture model fit to the prey swiming speed distributions are reported in Table S4 alongside their respective 95% confidence intervals.

For short trajectories, the prey swimming speed was positively correlated with trajectory length, but for trajectories longer than approximately four seconds this relation completely disappeared (see local polynomial regression fit in Fig. S10a). When limited to trajectories with lengths between four seconds and twelve seconds, the distribution of prey swimming speed still showed the same difference between the mixed and the monoculture (i.e. more prey individuals showed the faster swimming behavior in the mixed culture, compare Fig. S10b with Fig. S9a)

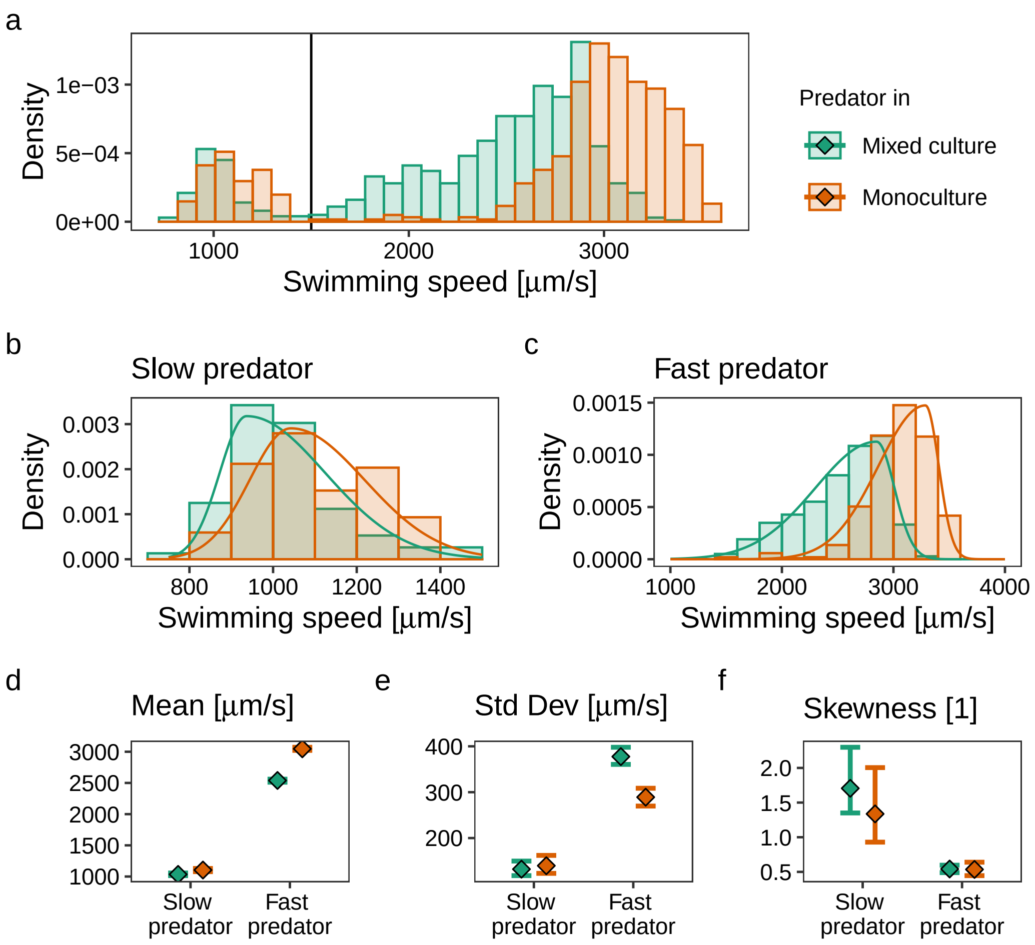


**Fig. S6** **a** Empirical density of the estimated swimming speeds of the predator individuals in the two different cultures. The vertical black line indicates the cut-off between slow- and fast-moving predators at $1500\,\mu\mathrm{m\,s^{-1}}$. **b** – **c** Empirical and fitted skew normal (black lines) densities for the swimming speed of the predator individuals in the two different cultures, respectively for the slow (**b**) and the fast moving (**c**) predators. **d** – **f** The estimated means (**d**), standard deviations (**e**) and skewness parameters (**f**) of the fitted skew normal distributions, for the fast and the slow predator in the two different cultures. The error bars indicate 95% confidence intervals

The difference in the prey swimming speed distribution between the mixed and the monoculture remained (or even increased) when for the mixed culture only the two videos were considered in which the prey abundance was comparable with the recorded abundances in the monoculture (see Fig. S2), as shown in Fig. S11 (compare with Fig. S9a).

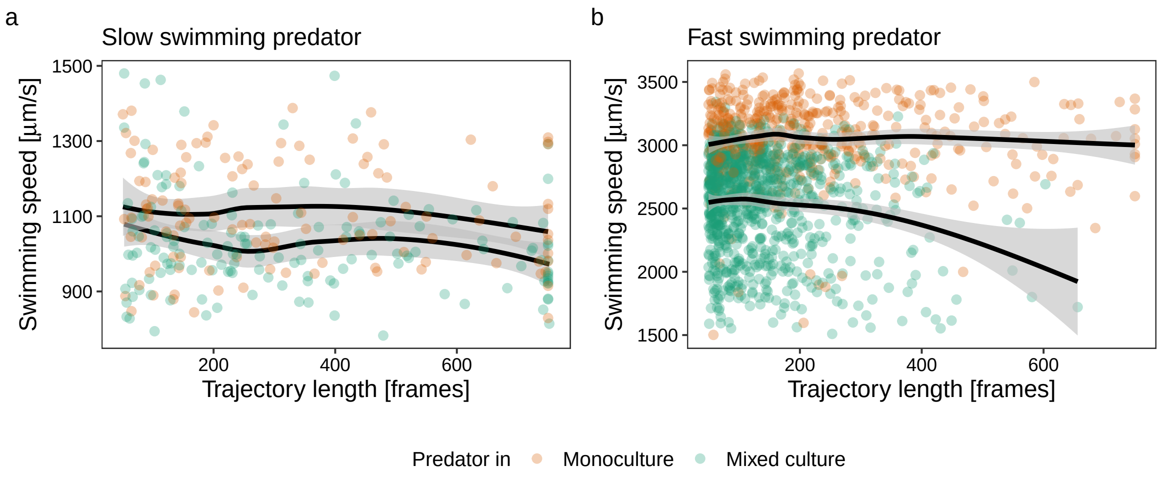


**Fig. S7** The predator swimming speed as a function of the trajectory length, colored based on culture. Each point represents a trajectory. The fits (solid lines) of loess local polynomial regressions are shown, with the shaded regions indicating 95% confidence intervals. **a** For the slow predator. **b** For the fast predator

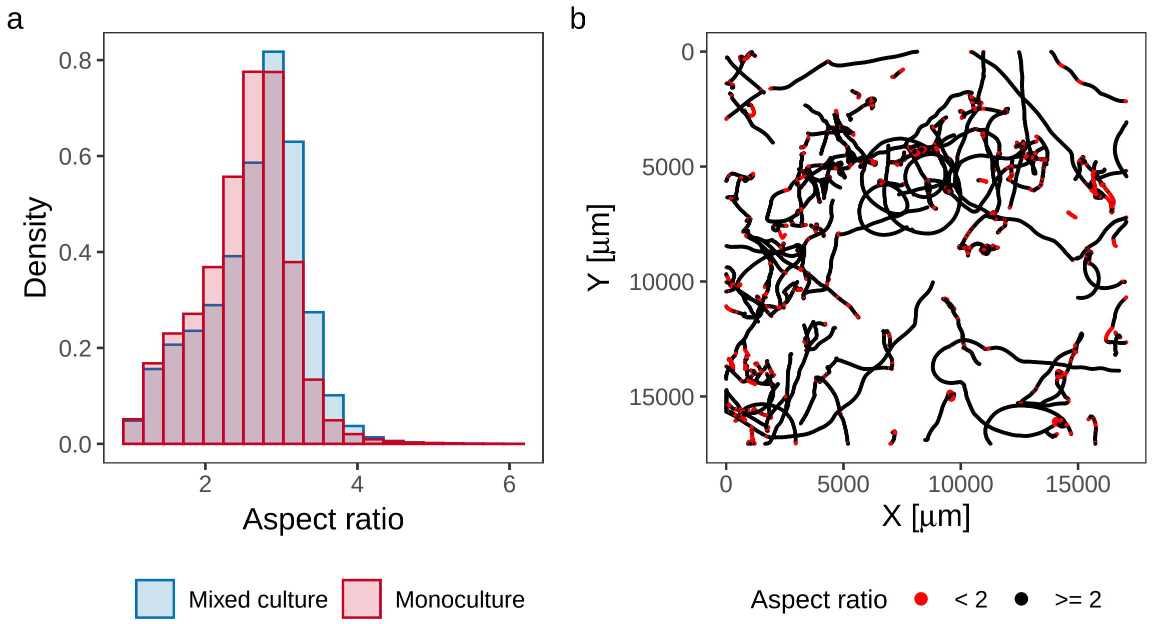


**Fig. S8 a**. The distribution as histograms of the aspect ratio of the prey individuals in both cultures. **b**. An example of the trajectories of a prey monoculture video. Each trajectory point at each frame is color-coded based on whether the aspect ratio of the predator individual was larger or smaller than two at that time point

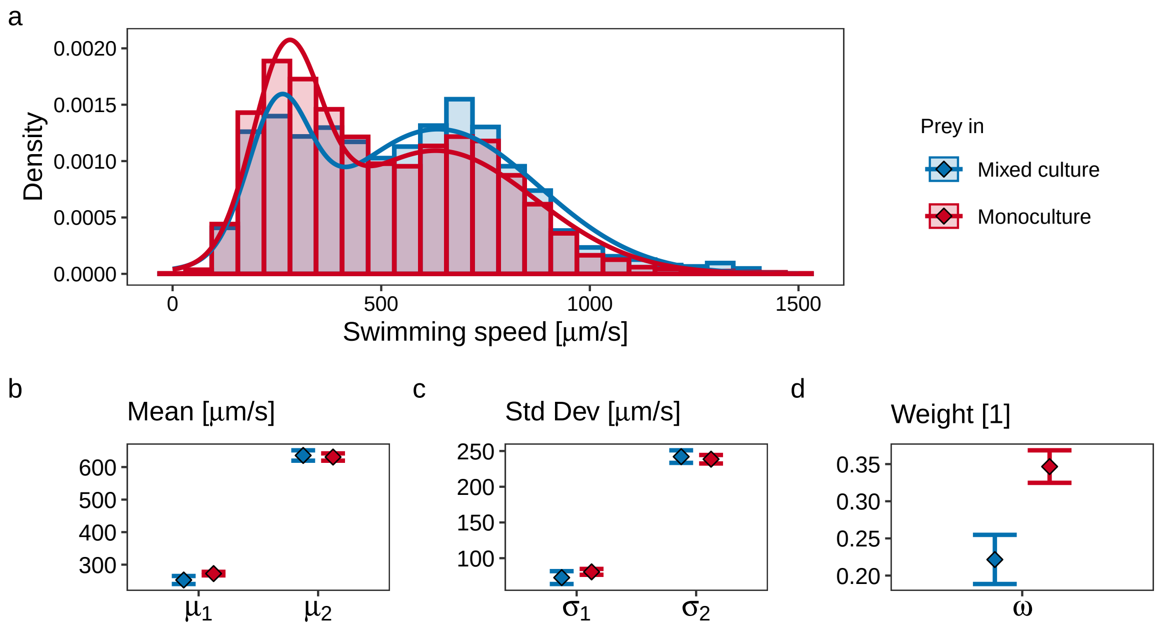


**Fig. S9** **a** Empirical and fitted Gaussian mixture (lines) densities for the estimated swimming speeds of the prey individuals in the two different cultures. **b** – **d** Respectively the estimated means, standard deviations and weights in the two different culture conditions. The error bars indicate 95% confidence intervals

**Table S4** Maximum likelihood estimates of the means, the standard deviations and the weights of the swimming speed distribution of the prey based on a weighted composition of two normal distributions in both the mixed and the monoculture

| Culture | Parameter | Estimate | 95% CI |
|---|---|---|---|
| Mono | $\omega$ | 0.347 | (0.32, 0.37) |
| | $\mu_1$ | 272.558 | (267.14, 277.92) |
| | $\sigma_1$ | 80.809 | (76.75, 84.95) |
| | $\mu_2$ | 630.796 | (619.65, 642.03) |
| | $\sigma_2$ | 238.571 | (232.55, 244.45) |
| Mixed | $\omega$ | 0.222 | (0.19, 0.25) |
| | $\mu_1$ | 252.840 | (240.55, 265.16) |
| | $\sigma_1$ | 72.765 | (63.83, 81.82) |
| | $\mu_2$ | 635.662 | (619.64, 651.49) |
| | $\sigma_2$ | 242.064 | (233.43, 251.00) |

#### S3 Predator displacements

The slow predator made significantly larger displacements in the monoculture (non-overlapping 95% confidence intervals, see Fig. 3 in the main text). This can also be seen in Table S5 where the mean square displacements are reported for time lags of one, five and nine seconds. This was also the case for the fast predator, but the confidence intervals were overlapping for the displacements after five and nine seconds. This overlap is likely to be caused by the fewer

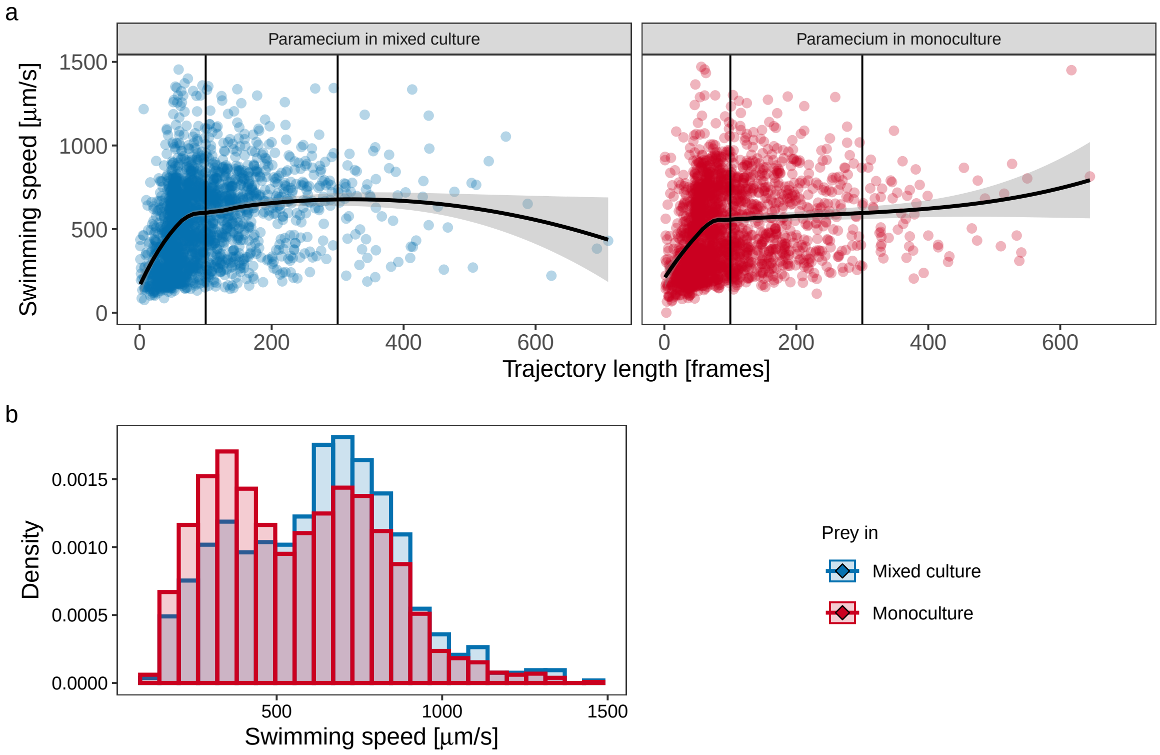


**Fig. S10 a** Prey swimming speed as a function of trajectory length, colored based on culture (left: mixed culture; right: monoculture). Each point represents a trajectory. The fits (solid lines) of loess local polynomial regressions are shown, with the shaded regions indicating 95% confidence intervals. The vertical blakc lines respectively denote the 100 and the 300 frames mark. **b** Empirical distributions for the estimated swimming speeds of the prey individuals in the two different cultures when only trajectories are considered that were between 100 frames (four seconds) and 300 frames (12 seconds) long

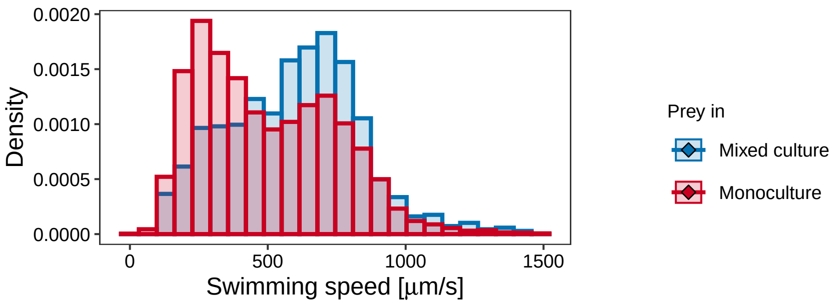


**Fig. S11** Empirical distributions for the estimated swimming speeds of the prey individuals in the two different cultures when only the two videos were considered in which the prey abundance was comparable with the recorded abundances in the monoculture

long trajectories for the fast predator (because the fast predator was quicker in swimming out of the captured area) which resulted in greater uncertainty in the estimated mean square displacements for longer time lags.

**Table S5** The estimated mean square displacements (and corresponding 95% bootstrapped confidence intervals) of the fast and the slow predator, respectively in the mixed and the monocultures and for three different time lags: one, five and nine seconds

| Time [s] | Culture | Movement | MSD [$mm^2$] | 95% CI |
|---|---|---|---|---|
| 1 | Slow | Mixed | 0.25 | (0.23, 0.27) |
| | | Mono | 0.34 | (0.31, 0.38) |
| | Fast | Mixed | 1.08 | (1.04, 1.11) |
| | | Mono | 1.23 | (1.17, 1.29) |
| 5 | Slow | Mixed | 3.64 | (3.15, 4.20) |
| | | Mono | 4.90 | (4.24, 5.59) |
| | Fast | Mixed | 13.74 | (12.72, 14.81) |
| | | Mono | 14.84 | (13.65, 16.11) |
| 9 | Slow | Mixed | 8.11 | (6.74, 9.67) |
| | | Mono | 12.66 | (10.25, 15.33) |
| | Fast | Mixed | 27.77 | (22.26, 34.66) |
| | | Mono | 30.33 | (25.93, 35.11) |

The estimated coefficients regarding the linear regression used to test the relation between the number of neighbors and the predator displacements are listed in Table S6 alongside their standard error and $p$-values. The table reports also the sensitivity analysis in which in addition to the detection range of $1500\,\mu m$, the detection ranges $1000\,\mu m$ and $2000\,\mu m$ were used to see how they influence the association between the number of neighbors and the predator displacements. The linear models were carried out separately for the fast and the slow predator.

Both the absence respectively the presence of prey and the number of local neighbors influenced the predator displacements (see Figs. 3 and 4). To investigate whether the presence or absence of prey or the number of neighbors was more important for the behavior of the predator, the predator swimming speed was used as the response variable in a linear regression. The mean number of local neighbors was an explanatory variables, alongside the culture (mixed or monoculture) and their interaction. The regression was carried out separately for the fast and the slow predator. The analysis of predator swimming speed showed that the predator swam faster in the monoculture than in the mixed culture regardless of number of neighbors, suggesting that the presence or absence of prey had a larger impact on predator swimming speed than the number of neighbors (Fig. S12 and Table S7).

**Table S6** The estimated coefficients and the corresponding standard errors of the covariates used in the linear regression with predator displacement as dependent variable. The *t*- and *p*-values according to the estimates and their standard errors are reported as well. For both the slow and the fast swimming predator, the regression model was fitted three times, once per different detection range. This served as a sensitivity analysis for the effect of the detection range

| Range | Covariate | Coefficient | Std. error | *t*-value | *p*-value |
|---|---|---|---|---|---|
| **Slow predator** | | | | | |
| 1000$\mu$m | Intercept | 475.22 | 6.04 | 78.72 | <0.0001 |
| | No. of prey neighbors | -22.13 | 10.65 | -2.08 | 0.0379 |
| | Predator monoculture | 58.77 | 8.90 | 6.60 | <0.0001 |
| | No. of predator neighbors | -13.28 | 13.09 | -1.01 | 0.3105 |
| | Predator monoculture × No. of predator neighbors | 22.94 | 20.73 | 1.11 | 0.2686 |
| 1500$\mu$m | Intercept | 480.42 | 4.32 | 111.21 | <0.0001 |
| | No. of prey neighbors | -17.61 | 5.36 | -3.28 | 0.0010 |
| | Predator monoculture | 56.33 | 8.37 | 6.73 | <0.0001 |
| | No. of predator neighbors | -9.34 | 7.03 | -1.33 | 0.1841 |
| | Predator monoculture × No. of predator neighbors | 21.50 | 12.41 | 1.73 | 0.0832 |
| 2000$\mu$m | Intercept | 495.27 | 8.20 | 60.42 | <0.0001 |
| | No. of prey neighbors | -14.68 | 4.46 | -3.29 | 0.0010 |
| | Predator monoculture | 30.01 | 11.69 | 2.57 | 0.0104 |
| | No. of predator neighbors | -12.04 | 5.11 | -2.36 | 0.0185 |
| | Predator monoculture × No. of predator neighbors | 29.55 | 9.29 | 3.18 | 0.0015 |
| **Fast predator** | | | | | |
| 1000$\mu$m | Intercept | 968.57 | 7.48 | 129.51 | <0.0001 |
| | No. of prey neighbors | -37.72 | 14.51 | -2.60 | 0.0093 |
| | Predator monoculture | 65.33 | 10.27 | 6.36 | <0.0001 |
| | No. of predator neighbors | -7.30 | 14.00 | -0.52 | 0.6020 |
| | Predator monoculture × No. of predator neighbors | -137.96 | 25.23 | -5.47 | <0.0001 |
| 1500$\mu$m | Intercept | 972.10 | 5.49 | 177.11 | <0.0001 |
| | No. of prey neighbors | -22.64 | 8.37 | -2.70 | 0.0069 |
| | Predator monoculture | 77.68 | 9.53 | 8.15 | <0.0001 |
| | No. of predator neighbors | -7.31 | 7.59 | -0.96 | 0.3354 |
| | Predator monoculture × No. of predator neighbors | -97.68 | 15.35 | -6.36 | <0.0001 |
| 2000$\mu$m | Intercept | 989.63 | 10.89 | 90.87 | <0.0001 |
| | No. of prey neighbors | -20.17 | 6.55 | -3.08 | 0.0021 |
| | Predator monoculture | 77.10 | 14.28 | 5.40 | <0.0001 |
| | No. of predator neighbors | -3.30 | 6.12 | -0.54 | 0.5902 |
| | Predator monoculture × No. of predator neighbors | -81.14 | 12.35 | -6.57 | <0.0001 |

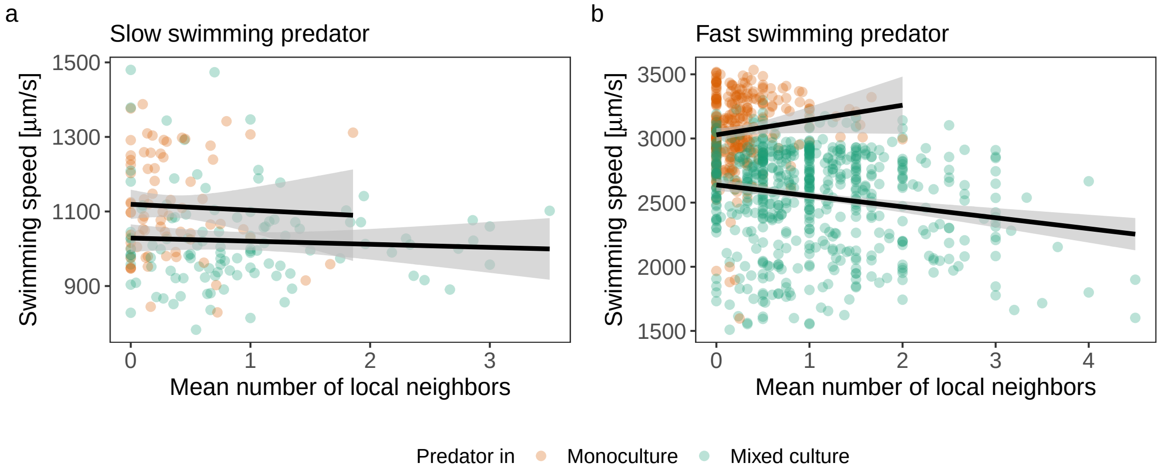


**Fig. S12** The predator swimming speed in the two cultures as a function of the mean number of local neighbors. The results of the linear regression and the corresponding 95% confidence intervals are included. **a** Slow-swimming predator. **b** Fast-swimming predator

**Table S7** Results of the linear regression in which the predator swimming speed (dependent variable) and the mean number of neighbors (independent variable) were used. The table lists the estimated coefficients and their standard errors alongside the the corresponding *t*- and *p*-values. The linear regression was fitted separately for the slow and the fast predator

| Covariate | Coefficient | Std. error | *t*-value | *p*-value |
|---|---|---|---|---|
| **Slow predator** | | | | |
| Intercept | 1028.63 | 18.64 | 55.17 | <0.0001 |
| Mean no. of neighbors | -8.31 | 15.48 | -0.54 | 0.5918 |
| Predator monoculture | 90.57 | 27.26 | 3.32 | 0.0011 |
| Mean no. of neighbors × Predator monoculture | -7.41 | 42.45 | -0.17 | 0.8616 |
| **Fast predator** | | | | |
| Intercept | 2637.99 | 23.64 | 111.61 | <0.0001 |
| Mean no. of neighbors | -85.29 | 18.00 | -4.74 | <0.0001 |
| Predator monoculture | 390.49 | 35.35 | 11.05 | <0.0001 |
| Mean no. of neighbors × Predator monoculture | 200.60 | 66.57 | 3.01 | 0.0027 |

#### S4 Prey grouping

The simulated uniform neighbor distance UND, i.e. the nearest neighbor distance for equally distanced individuals (i.e. cells), is shown in Fig. S13 as a function of how many individuals there are in the considered wnindow (i.e. area).

An alternative way to analyze prey grouping behavior is described here. To test whether prey individuals form bigger

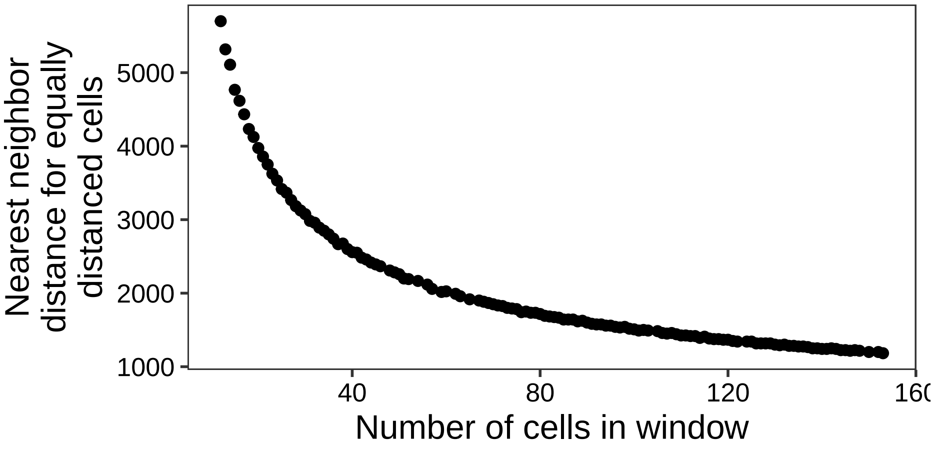


**Fig. S13** The nearest neighbor distance for equally distanced individuals (i.e. cells) as a function of how many cells there are in the window considered (i.e. the the area capture by the videos

groups and decrease the distance between other prey individuals every 25 frames (i.e., every second) a frame was selected from the videos of the prey monoculture and the mixed culture. Hence, as there were four replicates per culture and three 30 seconds long videos per replicate, 720 frames were selected. The area captured by the videos is the observation window $W$ for which the locations of the prey and the predator individuals are available. The locations of the prey individuals in each frame can be considered as a spatial point pattern and the respective intensity function $\lambda(u)$ is the expected number of individuals per unit length at any location $u$ inside the window $W$ for a given frame (Baddeley et al. 2015). An analysis based on the intensity functions of these spatial point patterns was carried out, build on the idea that more grouped individuals will produce higher intensities than more uniformly or randomly spread individuals. In other words, if the prey grouped significantly more in the presence of the predator, this difference should be quantifiable based on estimated intensity functions.

The intensity function $\lambda(u)$ of the observed spatial point pattern in each considered frame was estimated nonparametrically by kernel estimation using Diggle's estimator is (Diggle 1985) $\hat{\lambda}(u) = \sum_{k=1}^{n} \frac{1}{e(z_i)} \kappa(u - z_k)$, where $z_k$ ($k = 1, \ldots, n$) are the locations of the $n$ prey individuals in the considered frame (i.e. the spatial point pattern). Further, $\kappa(u)$ is the kernel function (here chosen to be a Gaussian density) and $e(u) = \int_W \kappa(u - v)\mathrm{d}v$ is a correction for edge effects.

As the number of prey individuals present in different frames and videos varied, the estimated intensity functions were further normalized. The integral of Diggle's estimator $\hat{\lambda}$ over $W$ is exactly equal to the observed number of locations (Baddeley et al. 2015); hence to normalize the estimated intensity functions it is sufficient to divide them by the number of prey individuals observed in the respective frames.

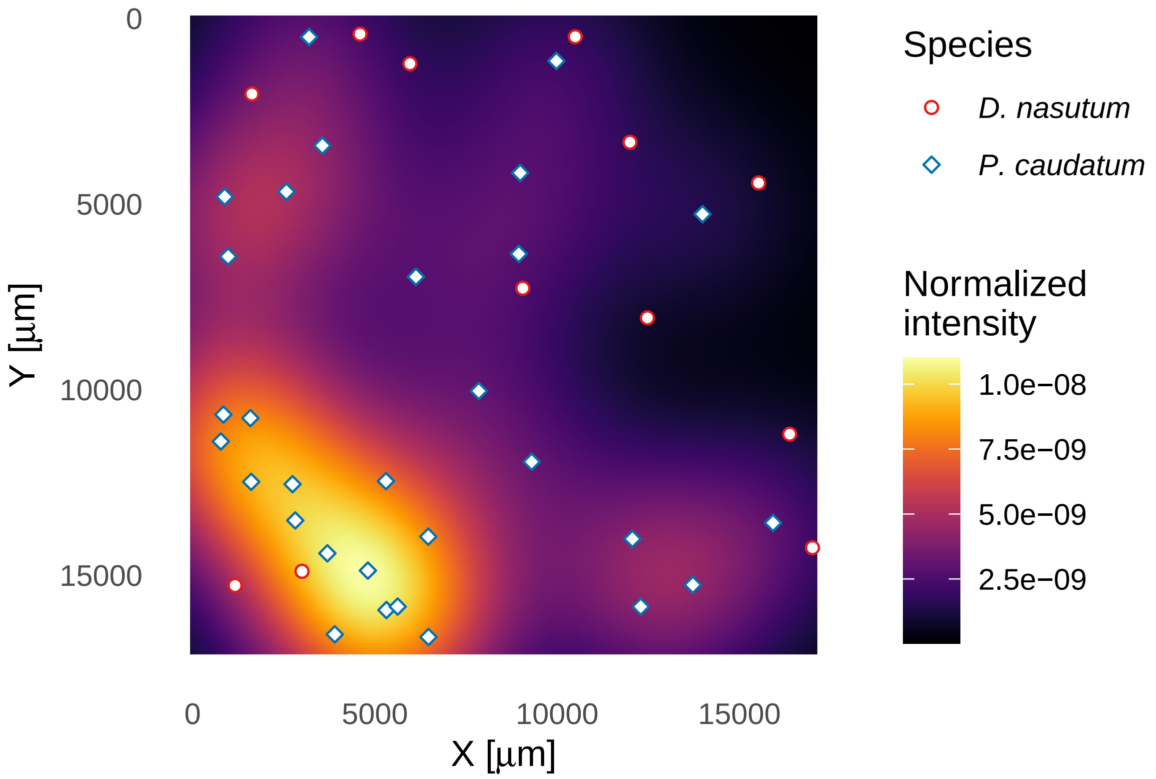


**Fig. S14** Example of a normalized estimated intensity function of a spatial point pattern consisting of locations of the prey individuals. The blue diamonds represent the prey individuals and the red circles are the predator individuals

Diggle's estimator $\hat{\lambda}$ is implemented in the function `density.ppp()` in the R package `spatstat` (Baddeley et al. 2015). The used function returns the estimated intensity function as a $128 \times 128$ matrix. Each estimated intensity function was then squared and summed up; hence, a sums of square measure was computed. Note that intensity functions that include regions with high intensities lead to bigger sums of square values than more uniform intensity functions.

An example for an estimated intensity function for a frame in a video is provided in Fig. S14, which also includes the locations of both the prey and the predator individuals. As can be seen, in this instance there is a region of higher intensity in the bottom left corner, where the prey grouped more. In this example, the predator individuals seem to be mostly in regions of lower intensities (i.e., with fewer prey individuals present).

To quantify the difference in intensity functions between the cultures, a simple mixed effects model was attempted. For this, the calculated sums of squares were log-transformed to ensure numerical stability (because the values had very small order of magnitudes, i.e., $10^{-13}$) and then used as the response variable in said mixed effects model. In

**Table S8** The estimated coefficients and their significances for the mixed models assessing prey grouping, both for the model based on distances and the model based on intensities

| Based on | Covariate | Type | Coefficient | Std. Err. | Test | Value | $p$-value |
|---|---|---|---|---|---|---|---|
| Distances | Intercept | Fixed | 0.4618 | 0.0083 | $t$ | 55.7833 | <0.0001 |
| | Mono | Fixed | -0.0334 | 0.0074 | $t$ | -4.5386 | <0.0001 |
| | Time | Fixed | -0.0003 | 0.0003 | $t$ | -1.0899 | 0.2761 |
| | Mono:Time | Fixed | 0.0004 | 0.0004 | $t$ | 0.9384 | 0.3484 |
| | Video:Replicate | Std. Dev. | 0.0186 | - | $\chi^2$ | 44.2674 | <0.0001 |
| | Replicate | Std. Dev. | 0.0071 | - | $\chi^2$ | 0.1272 | 0.7214 |
| | Residual | Std. Dev. | 0.0506 | - | - | - | - |
| Intensities | Intercept | Fixed | -28.9646 | 0.0203 | $t$ | -1428.9029 | <0.001 |
| | Mono | Fixed | -0.0857 | 0.0125 | $t$ | -6.8668 | <0.001 |
| | Time | Fixed | -0.0016 | 0.0005 | $t$ | -3.0713 | 0.002 |
| | Mono:Time | Fixed | 0.0025 | 0.0007 | $t$ | 3.3278 | <0.001 |
| | Video:Replicate | Std. Dev. | 0.0632 | - | $\chi^2$ | 240.1777 | <0.001 |
| | Replicate | Std. Dev. | 0.0000 | - | $\chi^2$ | 0.0000 | 1.000 |
| | Residual | Std. Dev. | 0.0858 | - | - | - | - |

this model, the culture composition was included as a factor with the levels "prey monoculture" and "mixed culture", the latter being the reference level. Further, time in seconds was included as a continuous covariate and the interaction between culture and time was included as well. Lastly, the video number was included as a random intercept nested in replicate. This is the same model that was used for the analysis of prey grouping based on the calculated Grouping Index (see main text).

The estimated parameters and their significances are reported in Table S8 for both the approach based on distances (see main text) as well as for the one based on intensities (i.e. the one described here). The approach for prey grouping based on intensities led to very similar results as the approach based on distances (compare Fig. 5 with Fig. S15), but in the case of the analysis based on the estimated intensity functions the variability in time was even larger which led to larger uncertainty in the results. Nevertheless, as two different approaches came to very similar results, this adds slightly to the evidence that there could be increased prey grouping in the presence of predator, although it could also be because of changes in prey density. In other words, the evidence is weak and the results inconclusive.

Lastly, Fig. S16a is the same as Fig. 5a but with a highlight on the two mixed culture videos in which the prey abundances were comparable to the abundances in the prey monoculture (see Fig. S2). As can be seen, in this figure there is no difference in prey grouping between the mixed and monoculture, which is also shown in Fig. S16b. This suggest that any difference in prey grouping between the mixed and the monoculture might be because of changes in prey densities and not because of the presence or absence of predators.

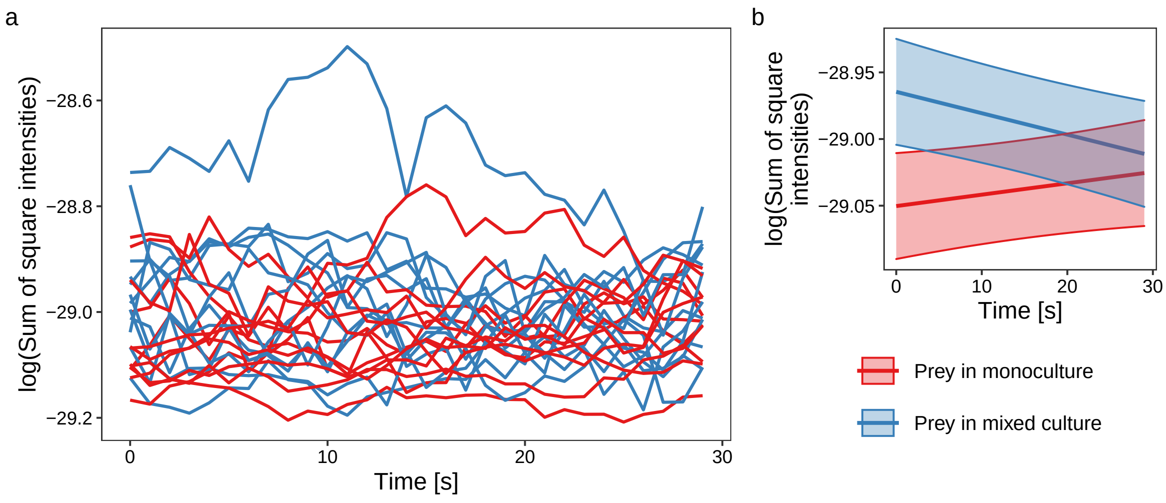


**Fig. S15** Prey grouping based on the estimated and summarized intensity functions with time in seconds on the x axis and colored based on culture. **a** The calculated data, grouped by videos (lines). **b** Results of the mixed effects models fitted. The 95% confidence intervals are based on fixed effects only

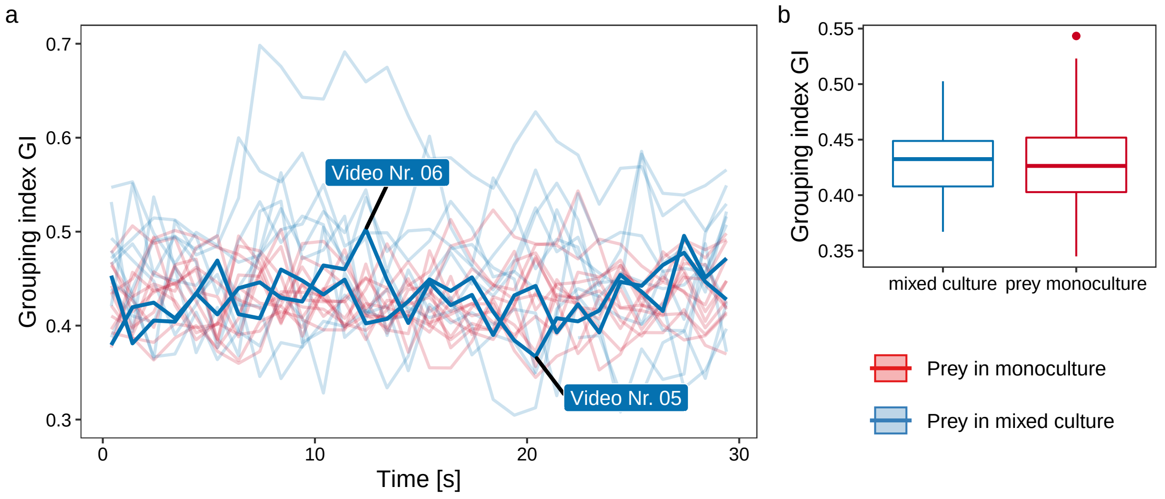


**Fig. S16** Prey grouping factor as a function of culture type and time, in the case of the mixed culture with focus on the two mixed culture videos in which the prey abundances were comparable to the abundances in the prey monoculture. **a** The calculated data, grouped by videos (lines). **b** Boxplots for the prey grouping factor for the two different cultures

#### S5 Morphology and speed

The estimates for the mean body size analysis and their confidence intervals are listed in Table S9. The body size of the fast swimming predator significantly decreased by 15.1% in the monoculture. This was not the case for the slow predator, where the decrease by 4.0% in body size in the monoculture was not significantly different from 0. The prey

**Table S9** Maximum likelihood estimates of the mean, the standard deviation and the skewness for the species body size distribution distribution of the predator based on a skew normal distribution. The estimates are given for both the slow and the fast moving predator as well as for the prey in both the mixed and the monoculture

| Species | Movement | Culture | Parameter | Estimate | 95% CI |
|---|---|---|---|---|---|
| Predator | Slow | Mono | Mean | 9893.11 | (9650.52, 10113.75) |
| | | | Std. Dev. | 1308.77 | (1147.55, 1479.71) |
| | | | Skewness | 0.81 | (0.61, 1.09) |
| | | Mixed | Mean | 10300.00 | (10051.56, 10561.10) |
| | | | Std. Dev. | 1600.00 | (1427.46, 1787.77) |
| | | | Skewness | 1.05 | (0.86, 1.32) |
| | Fast | Mono | Mean | 11669.98 | (11442.98, 11818.93) |
| | | | Std. Dev. | 2200.00 | (2068.92, 2346.53) |
| | | | Skewness | 1.59 | (1.34, 1.94) |
| | | Mixed | Mean | 13739.01 | (13621.83, 13877.40) |
| | | | Std. Dev. | 1960.37 | (1879.48, 2059.96) |
| | | | Skewness | 1.22 | (1.13, 1.33) |
| Prey | - | Mono | Mean | 5771.65 | (5736.62, 5816.27) |
| | | | Std. Dev. | 1692.57 | (1669.21, 1726.31) |
| | | | Skewness | 1.06 | (1.03, 1.10) |
| | | Mixed | Mean | 5726.98 | (5653.64, 5780.94) |
| | | | Std. Dev. | 1670.02 | (1629.56, 1721.92) |
| | | | Skewness | 1.18 | (1.12, 1.25) |

body size was not significantly different in the two cultures.

Fig. 2 shows the relation between species body size and species swimming speed. A linear model with swimming speed as the response variable and body size (named area in the model) and culture as covariates (with interaction) was fitted separately for the slow and the fast predator and for the prey. In the case of the fast predator there was no relation between body size and swimming speed, while for the slow predator and the prey there was a clear trend of swiming speed increasing with body size (Table S10).

**Table S10** Coefficients estimates for the covariates used in the linear regression with species swimming speed as the reponse variable, reported separatedly for the slow and the fast predator and the prey. The covariates were area (i.e. species body size) and culture (with levels mono and mixed), without interaction. Also reported are the standerd error, the test statistic and the corresponding *p*-values

| Species | Movement | Parameter | Estimate | Std. Err. | *t*-value | *p*-value |
|---|---|---|---|---|---|---|
| Predator | Slow | Intercept | 579.656 | 64.745 | 8.953 | <0.001 |
| | | Area | 0.044 | 0.006 | 7.042 | <0.001 |
| | | Monoculture | 93.915 | 107.841 | 0.871 | 0.385 |
| | | Interaction | 0.000 | 0.011 | 0.002 | 0.999 |
| | Fast | Intercept | 2503.792 | 82.557 | 30.328 | <0.001 |
| | | Area | 0.003 | 0.006 | 0.451 | 0.652 |
| | | Monoculture | 676.714 | 118.174 | 5.726 | <0.001 |
| | | Interaction | -0.014 | 0.009 | -1.519 | 0.129 |
| Prey | - | Intercept | 24.372 | 15.049 | 1.619 | 0.105 |
| | | Area | 0.092 | 0.003 | 36.455 | <0.001 |
| | | Monoculture | -4.946 | 17.675 | -0.280 | 0.780 |
| | | Interaction | -0.008 | 0.003 | -2.638 | 0.008 |
